# Carbon-phosphorous exchange rate constrains growth of arbuscular mycorrhizal fungal networks

**DOI:** 10.1101/2025.05.12.653230

**Authors:** Corentin Bisot, Loreto Oyarte Galveza, Felix Kahane, Marije van Son, Bianca Turcu, Rob Broekman, Kai-Kai Lin, Paco Bontenbal, Max Kerr Winter, Vasilis Kokkoris, Stuart A. West, Christophe Godin, E. Toby Kiers, Thomas S. Shimizu

**Affiliations:** Amsterdam Institute for Life and Environment, Vrije Universiteit, Amsterdam, The Netherlands; AMOLF Institute, Amsterdam, The Netherlands; Laboratoire Reproduction et Développement des Plantes, Univ Lyon, ENS de Lyon, UCB Lyon 1, CNRS, INRAE, INRIA, Lyon, France; Society for the Protection of Underground Networks (SPUN), Dover, DE, United States of America; Department of Biology, Oxford University, United Kingdom

## Abstract

Symbiotic nutrient exchange between arbuscular mycorrhizal (AM) fungi and their host plants varies widely depending on their physical, chemical, and biological environment. Yet dissecting this context dependency remains challenging because we lack methods for tracking nutrients such as carbon (C) and phosphorus (P). Here, we developed a new approach to quantitatively estimate C and P fluxes in the AM symbiosis from comprehensive network morphology quantification, achieved by robotic imaging and machine learning based on roughly 100 million hyphal shape measurements. We found that rates of C transfer from the plant and P transfer from the fungus were, on average, related proportionally to one another. This ratio was nearly invariant across AM fungal strains despite contrasting growth phenotypes, but was strongly affected by plant host genotype. Fungal phenotype distributions were bounded by a Pareto front with a shape favoring specialization in an exploration-exploitation trade-off. This means AM fungi can be fast range expanders or fast resource extractors, but not both. Manipulating the C/P exchange rate by swapping the plant host genotype shifted this Pareto front, indicating that the exchange rate constrains possible AM fungal growth strategies. We show by mathematical modeling how AM fungal growth at fixed exchange rate leads to qualitatively different symbiotic outcomes depending on fungal traits and nutrient availability.

## Introduction

The symbiotic partnership between arbuscular mycorrhizal (AM) fungi and their host plants is one of the most ubiquitous symbioses on Earth, with 70% of all plant species forming these resource exchange partnerships. Plants provide AM fungi with carbon in the form of sugars and fats, while phosphorous (P) and other nutrients are provided by fungal partners. In some cases, fungi have been shown to provide up to 80% of a host plant’s P supply (Smith and Read, 2010). In return, plants deliver on average 6% of their photosynthetically fixed carbon (C) to AM fungal partners (Hawkins et al., 2023). This partnership helped facilitate the colonization of land by plants over 400 million years ago, and today it is fundamental to both global ecosystem productivity and the regulation of the Earth’s climate(Smith and Read, 2010).

While the importance of nutrient trade between AM fungi and plants is well recognized, resource exchange is highly context dependent. Biotic and abiotic conditions influence the trade of nutrients in ways that we do not fully understand. This context dependency makes it difficult to predict cumulative nutritional effects of the symbiosis, such as the total amount of phosphorus transferred by the fungal partner to its host over a given time. Specifically, research has shown that fungal strains differ in the amounts of phosphorus that they supply to their host plants (Pearson and Jakobsen, 1993; Smith et al., 2003; Koch et al., 2017). These differences have been assumed to be related to differences in fungal traits and/or fungal trading strategies over space and time (Thonar et al., 2011; Koch et al., 2006).

We anticipate that fungal strategies are under selection for multiple, potentially conflicting tasks. Limited resources can be invested in exploration, local densification, reproduction or feeding the hyphal surface microbiome (Kakouridis et al. (2024)). Such multi-objective optimizations under constraints often gives rise to performance tradeoffs identifiable in the form of Pareto fronts, boundaries in trait space that delineate achievable fungal phenotypes under constraints such as resource limitation Shoval et al. (2012). Resolving such Pareto fronts from experimental data can be informative of the type of constraints that underlie complex phenotypes such as morphogenesis.

Indeed, plants appear to adjust the amount of carbon they supply to fungi, in response to how much phosphorous they are supplied often referred to as reciprocal exchange (Jakobsen et al., 1992; Kiers et al., 2011; Hammer et al., 2011). However, the mechanisms that underlie such changes in nutrient exchange, and the strategies they might represent remain unclear (Bunn et al., 2024; Wyatt et al., 2014). Do plants and fungi have fixed trading rules, or do they modulate their trade behaviors depending on nutrient availability and/or the specific partners they interact with? And within a given plant-fungal pair, is the relative amount of nutrients received by each partner (*i*.*e*. the effective exchange rate) fixed in a manner that guarantees steady returns, or does it vary in a manner that might reflect more flexible strategies at play (e.g. cooperation vs cheating)? Past work has quantified the C/P-exchange ratio in relative terms using isotope labeling (Lekberg et al., 2024; Kiers et al., 2011). Yet, such end-point measurements are laborious and are often noisy due to the multiple extraction steps involved. Furthermore, they yield only a single ratio value per mycorrhizal replicate. An ideal experiment would allow tracking of both carbon and phosphorus transfer over time within each replicate. However, so far such dynamical *in vivo* tracking of two-way resource transfer has remained out of reach.

Here, we addressed this challenge by developing a novel approach based on high-throughput robotic imaging and machine learning that enables estimation of both carbon expenditure by the plant and phosphorous delivery by the fungus. Because the imaging measurements are non-invasive, we are able to obtain these estimates at regular (typically 2-hour) intervals as the mycorrhizal network develops. The key idea is that both the carbon cost of network growth (which dominates fungal carbon expenditure) and phosphorus uptake rate (which limits phosphorous transfer to the plant) can be estimated from precise measurements of hyphal morphology if such measurements can be conducted comprehensively throughout the *entire network* and during *steady-state growth*. The basic rationale is that both building costs and nutrient uptake rates of the hyphal network scale with morphological observables. First, the amount of carbon required for network construction is to a reasonable approximation proportional to the total volume *V* of the hyphal network (Bar-On et al., 2018). Second, the ability of the network to absorb solutes such as phosphorus from the environment and transfer them to the plant host is expected to depend on its total membrane surface area, *S*. We were able to quantify this dependence with direct measurements of phosphorus depletion and translocation to calibrate the dependence of phosphorous absorption and transfer to the host root on the network surface area *S*. Thus, sufficiently accurate measurements of morphology across the entire network, with appropriate calibrations, can enable reliable quantification of transferred carbon and phosphorous.

To precisely and comprehensively measure hyphal morphology across the entire network, we leveraged a robotic imaging and analysis pipeline we recently developed (Oyarte Galvez et al., 2025), capable of extracting the full network graph of the extraradical mycelium (ERM) of AM fungi over time, after forming symbiosis with an *in-vitro* root-organ-cultures (ROC) as host. In each of these samples, the colonized host was restricted to the root compartment, while the fungal network crossed a physical barrier to a second compartment (i.e. fungal side) lined with a permeable cellophane layer to optimize visualization. Given the approximately cylindrical geometry of hyphal filaments (Fig. 1A), the relevant morphological parameters are the hyphal radius *r* and total hyphal length *L*, which are related to surface area and volume by simple scaling relationships (*S* ∼ *rL* and *V* ∼ *r*^2^*L*). While the machine vision techniques developed in (Oyarte Galvez et al., 2025) sufficed to yield accurate measurements of the network length *L*, challenges remained in accurate determination of the hyphal radius *r*, given the relatively low magnification at which the robot images the hyphal network. Past estimates of AM hyphal radii vary over a broad range, from 0.6 to 9*µm* (a 15-fold difference)(Dodd et al., 2000), but whether that diversity reflects changes over time, variation across space, and/or differences between strains/species remained largely unknown. We therefore developed a new machine-learning based method for precisely extracting *r* for every hyphal edge within growing extraradical mycorrhizal (ERM) networks. This new method allowed us to obtain roughly one million hyphal radius measurements per network timelapse, and we applied it to ∼ 100 timelapses in this study.

**Figure 1.**
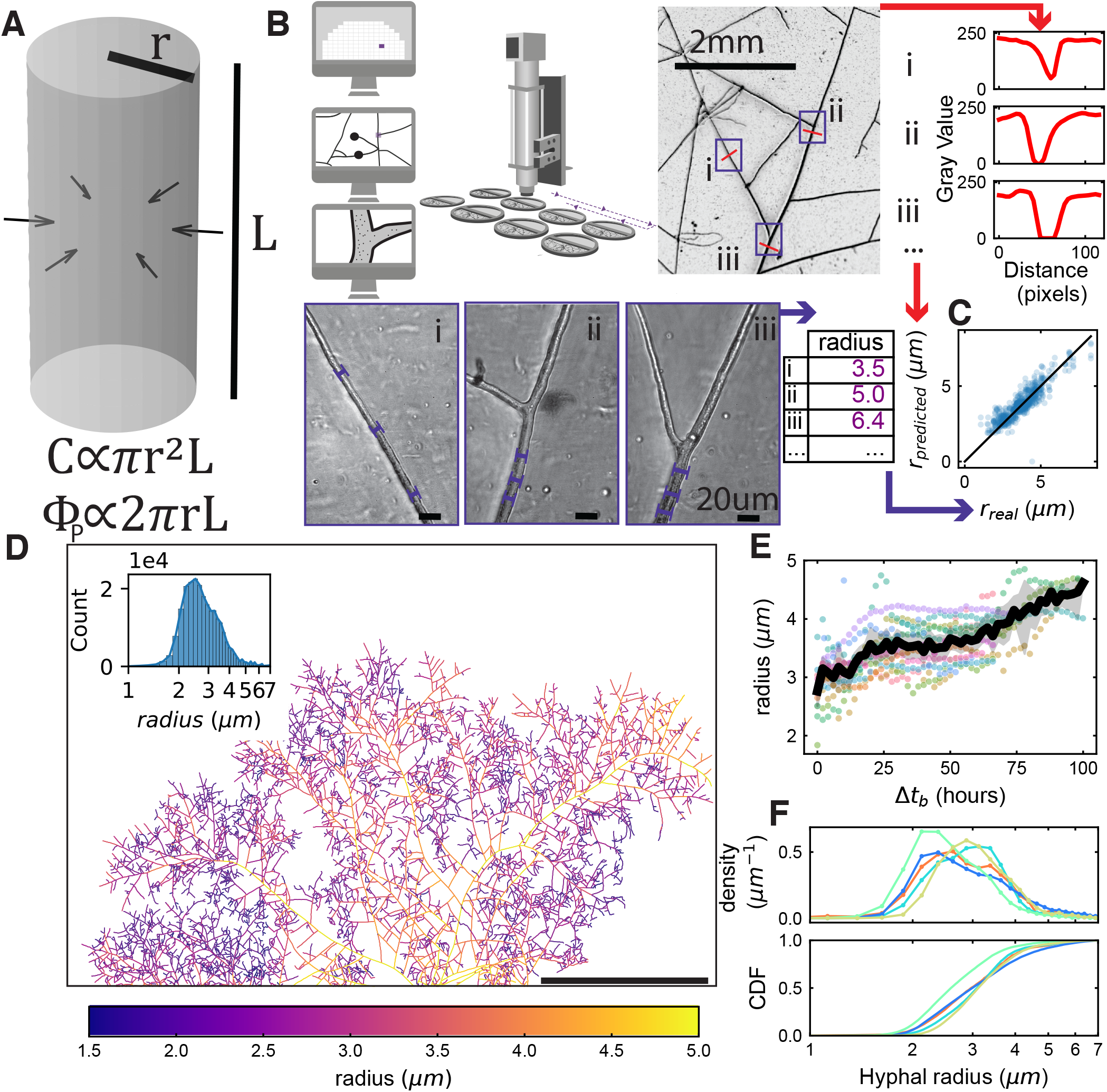
Hyphal radius variation in space, time and across genotypes as revealed by Convolutional Neural Network. **(A)** Illustration showing the importance of hyphal radius for fungal ressource economy. Cost of building a hyphal segment of length *L* and radius *r* is proportional to cylinder volume while absorption of solute from the environment is proportional to cylinder surface area. **(B)**Description of the dataset generation and training scheme. Images are acquired in low (scale bar is 2mm) and high magnification (scale bar is 20*µ*m) at different location in the network. Hyphal radius is manually measured on the high magnification images and orthogonal 120 px transect are extracted from the low magnification images. A Convolutional Neural Network is trained to predict the measured radius from the extracted transect. **(C)** Evaluation of the trained neural network on an independent dataset that has not been used for training (RMSE=0.70, R2 =0.66). Black line is the identity function, blue points are individual predictions from the independent dataset. **(D)** Network graph where each edge colour is mapped to the radius value estimated with the trained Convolutional Neural Network. Inset is the distribution of radius value over all 10px segments that constitute the graph. **(E)** Temporal variation in radius of a selection of (n=13) major hyphae of a network as a function of the time since their emergence after branching. Each dot colour correspond to a given hypha. Black line corresponds to average and grey shade to 95% confidence interval computed on all points over intervals of two hours. **(F)** *upper* Volume weighted distribution of the radius of all edges at final timestep for different fungal strains and species: R. *irregularis* A5 (cyan, n=8), C2 (blue, n=19) and C3 (red, n=4), R. *aggregatus* (green, n=4), G. *clarum* (yellow, n=4). Curves are linking the top of histogram bins. *lower* Cumulative distribution function representing for each radius the proportion of total network volume that is contributed by hyphal element below that radius.

For the validity of our morphology-based approach to estimating C/P exchange, it is important that measurements are made during steady-state growth, in which the cell’s elemental composition (Saldanha et al., 2004), as well as gene expression (Rautio et al., 2006; Pelechano and Pérez-Ortín, 2010) is constant. We ensured that the AM fungal network was in the steady-state growth regime by restricting our measurements to early times (*<* 200 h) after hyphal crossing into the fungal compartment. This is when the ERM network expands as a travelling-wave, with constant speed and density (Oyarte Galvez et al., 2025). Whereas carbon cost will be to a good approximation proportional to the network volume *V* under these conditions, the dependence of phosphorus transfer on surface area *S* requires some model of phosphorous uptake/transport. We therefore conducted direct measurements of phosphorus depletion and translocation to validate and calibrate a minimal model of phosphorous transport. Combined with this calibrated model, robotic imaging and analysis of network morphology allowed us to dynamically track how AM fungi utilize plant-derived carbon to acquire and transfer phosphorus to the plant. We quantified these dynamics for multiple combinations of plant × fungus × phosphorous concentration to investigate the statistics of C/P exchange across time and how this varied with changes in trade partners and environmental conditions. Remarkably, we found that despite considerable variation in C/P exchange across samples and across time, on average the amount of phosphorous transferred to the plant was approximately proportional to the amount of received carbon. Finally, we analyzed the significance of this proportional exchange for symbiotic trade using the framework of Pareto fronts, and extending a model of travelling-wave growth of AM fungal networks to account for C/P exchange.

### Machine-learning model reveals temporal, spatial and genetic variation in hyphal width

First, we asked if and how the absorbing capacity of AM fungal networks changed over time by measuring hyphal radii from three different AM fungal strains over a 5-6 day period. To obtain network-wide hyphal morphology measurements for estimating C expenditure and P absorption, we developed a machine-learning-based approach to extract hyphal widths from data generated by our high-throughput imaging robot (Fig. 1B).

We trained a Convolutional Neural Network (CNN) to extract the radius *r* of hyphal filaments from one-dimensional intensity linescans of hyphal transects within low-magnification output images of the robot (Fig. 1B). We selected approximately 1000 positions from 6 fungal networks and imaged them at both low (2x) and high (50x) magnification. To generate ground truth data for training, we first manually measured the hyphal radius *r* within all high resolution images. Subsequently, transect linescans orthogonal to the hyphal axis at those same positions were automatically extracted from corresponding low-magnification images. We then used the ground truth data as labels and the linescan data as features to train a two-layer CNN to extract *r*. We evaluated the CNN’s performance on an independent test set (RMSE=0.70, R2 =0.66), confirming that the model could reliably predict *r* from data outside the training set (Fig. 1C, Fig. S1).

The resulting model can be used to efficiently extract *r* for every hyphal edge of the fungal network (Fig. 1D). We found that the radius of hyphae *r* was distributed broadly (Fig. 1D Inset), from around 1 *µm* for the thinnest hyphae of branched absorbing structures (BAS; key sites for nutrient uptake) to about 6-7 *µm* for the thickest runner hyphae (RH). In addition to this spatial variation, we observed that RH tended to increase their width over time, with values of *r* growing, on average, from 3 *µm* to 4.5 *µm* over 100 h (Fig. 1E). Such increases were not due solely to cell wall thickening, because the cell wall never contributed to more than 20% of total hyphal width (Methods). This hyphal widening therefore represents increases in both plasma membrane surface area and cell volume. We also ruled out that these temporal changes were artifacts of the CNN model by confirming hyphal widening in manually measured subsets of edges imaged over multiple days at high magnification (Fig. S2). Fungal hyphae are generally believed to grow only axially, by elongation at their tips, thereafter maintaining an approximately constant hyphal width Steinberg et al. (2017); Riquelme et al. (2018) unless subjected to (e.g. osmotic) stresses (Mazheika et al., 2024). Evidently, AM hyphae also grow radially, by expanding the width of already built hyphal tubes over time.

Next, we asked whether the distribution of the hyphal radius *r* varied across AM fungal genotypes, focusing on *Rhizophagus irregularis* A5, C2 and C3, *R. aggregatus*, and *Rhizophagus clarum* (see Methods), using the CNN model to predict *r* for all network edges. The choice of genotypes was driven by the motivation to sample broadly across the variation in network growth phenotypes, and to ensure sufficient coverage across the range of fungal traits. These strains were able to be cultured with *in-vitro* culture. We found that the *r*-distribution for all tested genotypes were broad, with considerable overlap in the distributions across genotypes (Fig. 1F). Specifically, we found that *R. aggregatus* allocated proportionally less of its total network volume to thicker hyphae compared to *R. irregularis* (Fig. 1F Lower).

### Pattern of fungal growth suggests progressive relaxation of carbon constraint

To place these contrasting hyphal radius distributions in the context of carbon costs, we next quantified the spatial density of hyphal filaments *ρ* (*µm/mm*^2^), as well as the range-expansion speed *v*_wave_ of the network. This was inspired by a previous study in which we showed that AM fungal network growth is well-described as a travelling-wave, with the spatial range of the fungal network (regions of nonzero hyphal filament density) expanding at a constant speed *v*_wave_ (Oyarte Galvez et al., 2025). In the present study, quantification of the radius *r* allowed us to compute the volume of every hyphal edge of the network. We also detected all spores of the network as well as their radius which allowed us to estimate their volume. At later times, spores could indeed contribute 10-20% of total fungal biovolume. The sum of hyphal and spore volumes could in turn be used to compute the total amount of carbon *C* in a given region of area *A*, and hence also (on dividing by *A*) the spatial density of carbon *ρ*_*C*_ (≡ *C/A*). We found that fungal networks that expanded faster (*i*.*e*. with greater *v*_wave_) tended to grow sparser (*i*.*e*. at lower *ρ*_*C*_) and vice versa (Fig. 2A). To further investigate whether this negative correlation between speed and density was robust across different plan hosts, we measured *ρ*_*C*_ and *v*_wave_ for three of the fungal strains in combination with two different host genotypes. We chose two different genotypes of carrot *Daucus carota* as these hosts (hereafter referred to as genotype 1 and genotype 2) grow at different rates, with genotype 1 growing slow and genotype 2 growing fast (Oyarte Galvez et al., 2025). We found that while both *v*_wave_ and *ρ*_*C*_ were affected by host genotype, they changed in a manner that preserved the negative correlation (Fig. S3). Specifically, networks grown with genotype 2 carrot root expanded their spatial range faster than those grown with genotype 1 carrot root, but when attached to either host, *R. irregularis* A5 grew sparser and expanded faster than *R. irregularis* C2, itself faster and sparser than *R. aggregatus*.

**Figure 2.**
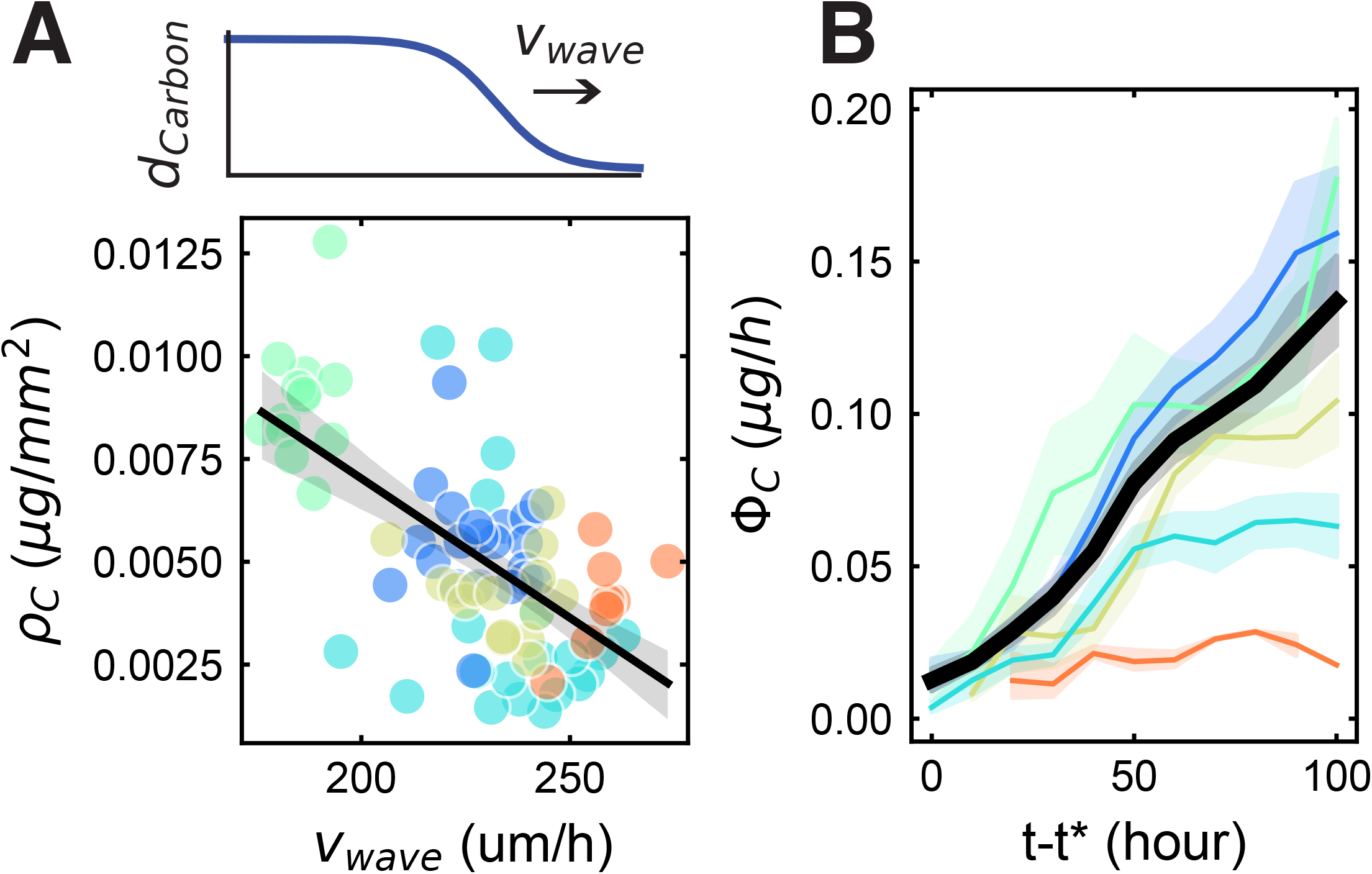
Carbon expenditure pattern of various fungal strains. **(A)** *top*: Illustration of typical fungal density wave dynamic. *bottom* Travelling-wave observables. *ρ*_*C*_ represents the instantaneous density of carbon across the network’s spatial range and *v*_wave_represents the instantaneous wave speed. Each small circle corresponds to an independent median of those parameters over 10 hours for all plates of the same strain (*n*= 7-11). Black line represent bootstrap regression on all datapoints. Grey shade correspond to 95% C.I. of the linear fit obtained via bootstrapping. **(B)** Carbon spent by the growing network in hyphal structures per hour as a function of time. Black line and shade represent average and 95% confidence interval of all replicates’ dynamics. Coloured lines correspond to average and 95% confidence over all replicates for one strain. Average and 95% confidence interval (mean *±*2*×* s.e.m.) are computed over 10-hour time intervals. For all plots, colour correspondence and number of replicates are the following: *R. irregularis* A5 (cyan, n=9), C2 (blue, n =19) and C3 (red, n=3), *R. aggregatus* (green, n=4), *G. clarum* (yellow, n= 4).

Both range expansion speed (promoting exploration) and hyphal density (promoting exploitation) are phenotypic traits of network growth. The existence of a negative correlation between growth traits therefore suggests the possibility of an exploration-exploitation trade-off in AM fungal growth strategy. Such a trade-off could arise from constraints due to a limiting nutrient resource. Unlike free-living filamentous fungi, AM fungi, as obligate biotrophs, rely entirely on their plant host for their carbon supply (Parniske, 2008). Given that the tradeoff itself was dependent on host identity, we suspected that host carbon and not other nutrients (Nitrogen, Phosphorous or other macro/micro nutrients) could be directly limiting. We therefore estimated how much carbon was expended by the fungal network on hyphal and spore growth at each timestep. Because the amount of carbon per unit volume of cells is known to be constant during steady-state growth (Bar-On et al., 2018), total carbon *C*_*t*_ in the steadily expanding network could be estimated by quantifying the total network volume *V* (Methods). We found that the rate of carbon expenditure Φ_*C*_, defined as the time derivative of total carbon in the network (Φ_*C*_ ≡ *dC*_t_*/dt*) increased substantially (by up to ∼ 10-fold) over time for all strains observed (Fig. 2B). Thus, if carbon is indeed the growth-limiting resource, this increase in Φ_*C*_ implies that carbon limitation of fungal growth is being progressively relaxed over time.

### Fungal P supply scales with network surface area

What could account for such a large increase in carbon input from the plant host during fungal network growth? Because AM fungi forage and exchange phosphorus to obtain plant carbon, we hypothesized that the evident increase in C supply from the plant host reflects a concomitant increase in P transfer to the plant root via the fungal network. To test this idea, we proceeded to quantify the amount of environmental P absorbed by the fungal network and transferred to the root. Using a spectrophotometric assay (see SI), we measured the total mass of phosphorus in the fungal compartment, the gel of the root compartment, and within the host root (Fig. 3A). Next, we estimated the total surface area of the network *S*(*t*) at each point in time until harvest using the network morphology extracted from our imaging pipeline (3B).

**Figure 3.**
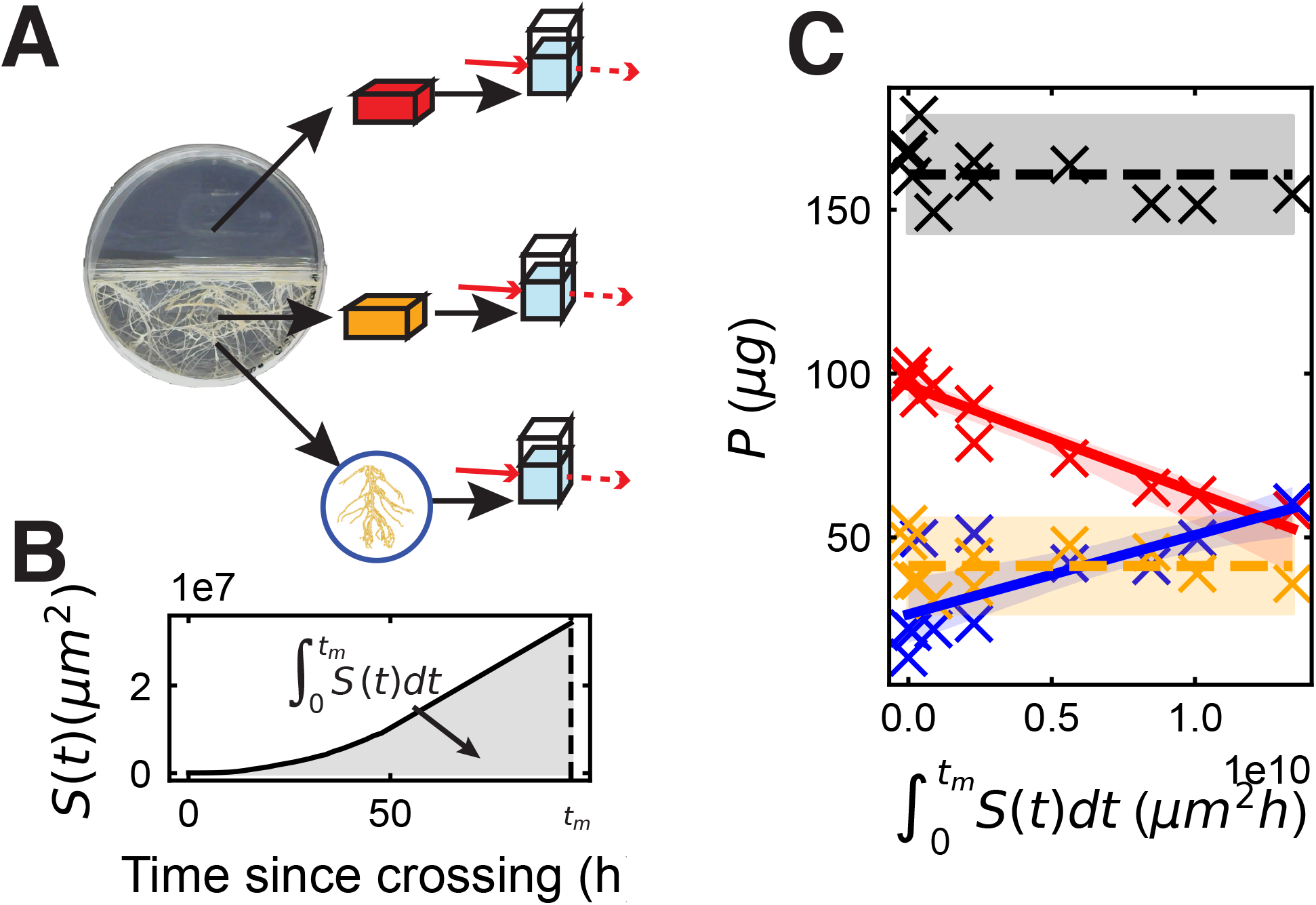
Phosphorous depletion and transfer by the growing network. **(A)** Illustration of the experimental method used for measurement of phosphorus mass in the fungal compartment (red), the gel of the root compartment (orange) and the host root (blue) **(B)** Network surface area *S*(*t*) (black curve) plotted up to harvest at time *t*_*m*_ for a typical plate. Grey shaded area under the curve highlights the time-integrated surface area, a key quantity in our analysis. **(C)** Total mass of phosphorous measured in the fungal compartment (*P*_f_, red), the gel of the root compartment (*P*_g_, orange) and the host root isolated from the root compartment (*P*_r_, blue) as a function of the time-integrated surface area. Black line and dots correspond to the total phosphorous *P*_t_ across all three measurements. Points correspond to individual replicates, full lines correspond to linear fit, dashed horizontal lines show averages for the corresponding category. Shades of full lines correspond to the 95% C.I. of the linear fit obtained via bootstrapping and shades of dashed lines to (mean ±2× s.e.m.). Coefficient of determination for the linear fits (*R*^2^) are 0.91 (red), 0.51 (blue) and Roose, 2006):

Assuming the absorption of P by the fungal network is limited by the number of P transporters on the hyphal surface at some characteristic density(Armstrong, 2008), the rate of P absorption Φ_*P*_ will be a product of the total surface area of the network *S*(*t*) and the rate of transport per unit area *J*. The depletion of total phosphorus in the fungal compartment *P*_f_ due to absorption can then be expressed as,

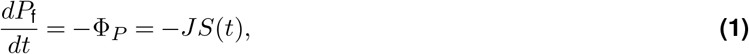

where the rate coefficient *J* may depend on the extracelullar concentration of P. Integrating Eq. 1 up to the measurement time *t*_m_ yields,

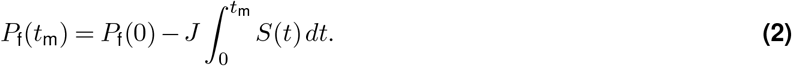

which predicts that the amount of P left in the compartment *P*_f_ will decrease linearly as a function of the time integrated surface area 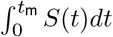 with slope −*J*. We found that the measured *P*_f_(*t*) did indeed decrease linearly with integrated surface area (Fig. 3C, red), and the slope allowed us to extract *J* ≈ 3 ng mm^−2^h^−1^. This transport parameter is important for the plant-fungal symbiosis because it determines the rate at which a given network will absorb P from its environment. For example, a network with this value of *J*, total length 1m, and an average radius 3 *µm* would extract about 1 *µg* of P per day. In contrast to *P*_f_, the amount of phosphorus *P*_g_ in the gel of the root compartment (which represents a baseline amount of inaccessible phosphorous, presumably adsorbed to the gel matrix (Jain et al., 2009)) did not significantly change throughout network growth (Fig. 3C, orange).

An expression similar to that in Eq. 1 had been proposed in previous models of P absorption by AM fungi (Schnepf

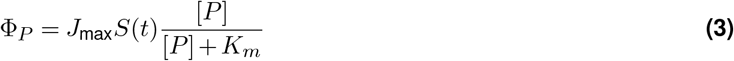

where [*P* ] represents the concentration of accessible phosphorous in solution within the fungal compartment *P* ≡ *P*_f_ − *P*_g_. Our experimentally validated formula (Eq. 1) corresponds to the saturated regime of Eq. 3 where the Michaelis-Menten constant *K*_*m*_ is small compared to the solute concentration (*i*.*e. K*_*m*_ ≪ [*P* ]) and hence *J* ≈ *J*_max_. The fact that *P*_f_ continues to decay linearly down to levels very close to the background level *P*_*b*_ suggests that the value of *K*_*m*_ is very small compared to [*P* ]_0_. Our estimate *J*_max_≈ *J* ≈ 3 ng·mm^−2^h^−1^ obtained by imaging network morphology and P depletion within *in vitro* networks is in good agreement with the range of values obtained in those previous studies (*J*_max_ ≈ 4−10 · ng mm^−2^h^−1^) based on fungal density and P concentration measurements in soil (Schnepf and Roose, 2006; Li et al., 1991). We also found that the decrease in *P*_f_ was balanced by a complementary increase in the amount of P in the host root *P*_r_ (Fig. 3C, blue). Given that neither the amount of phosphorus *P*_g_ in the gel of the root compartment (Fig. 3C, orange) nor the total amount of phosphosrus *P*_t_ (= *P*_f_ +*P*_r_ +*P*_g_) across all three measurements (Fig. 3C, black) significantly changed in the meanwhile, the increase in *P*_r_ can be wholly attributed to the symbiotic transfer of P across compartments via the fungal network and into the host root. Taken together, these data suggest that in these experiments, nearly all the P absorbed by the fungal network is efficiently transferred to the host root, without obstructions or delays that would result in a build up of P inside the fungal hyphae as had been observed in other studies (Kiers et al., 2011; Whiteside et al., 2019). This latter conclusion is further supported by considering our results in the context of literature reports of P concentration in AM fungal hyphae. The magnitudes of P absorption and network volume we measured suggest that approximately 50% of fungal dry biomass would be P if no P were transferred to the root (Fig. S4). Yet across the literature, the highest reported mass fraction of phosphorus in AM fungal hyphae was 4% of total dry weight (Hammer et al., 2011) while poly P content in yeast does not seem to greatly exceed 10% (Kornberg and Rao, 1999).

Taken together, we found through these experiments that (1) nearly all of the P absorbed by the network is transferred to the root without delay, and (2) the rate of P transfer from the fungal network to the host root is therefore equivalent to Φ_*P*_ . Consequently, the model of Eq. 3, calibrated with values of *J*_max_ and *K*_*m*_ obtained from our P depletion measurements, provides a means to estimate the rate of P transfer to the root from imaging-based measurements of total network surface area *S*(*t*).

### Proportionality between C expenditure and P supply reveals a plant-dependent exchange rate

Within our two-compartment petri dish setup, the fungus has access to C only through the plant host. Likewise, at the time where the fungal colony is being imaged, the plant host has access to P only through the fungal network. We therefore reasoned that examining the relationship between the rate of phosphorous transfer (Φ_*P*_ ) and the rate of carbon expenditure (Φ_*C*_) at every time point could provide insight into the nature of nutrient exchange. Despite considerable variation across replicates, we found that the relationship between these two rates averaged across all replicates (Fig. 4A, blue curve) was well-approximated by a straight line (Fig. 4A, black curve), with a slope of approximately 3 mass units of C per 1 mass unit of P. In fact, a constant ratio between Φ_*C*_ and Φ_*P*_ can be shown to be an expected consequence of travelling-wave growth in the long time limit, where the ratio is predicted to become proportional to the carbon density of the wave *ρ*_C_ behind the advancing front (specifically, Φ_*C*_*/*Φ_*P*_ → *ρ*_*C*_*/*[*P* ]_0_; see Methods). However, the data of Fig. 4A begins at times ( ≤ 5 days) much earlier than the onset of such a steady state ( ≈10 days). Furthermore, we found that the on-average linear dependence of Φ_*C*_ on Φ_*P*_ was recapitulated across all five tested AM fungal genotypes (Fig. 4B, S5) and, moreover, the slopes of these dependences were approximately the same (Fig. 4C) despite wide variation in carbon density of the wave *ρ*_*C*_ and its propagation speed *v*_wave_ (Fig. 2A). These data suggest that the observed proportionality of resource exchange is not a simple consequence of travelling-wave growth, but instead reflects the existence of one or more regulation mechanisms that ensures an approximately constant exchange rate.

**Figure 4.**
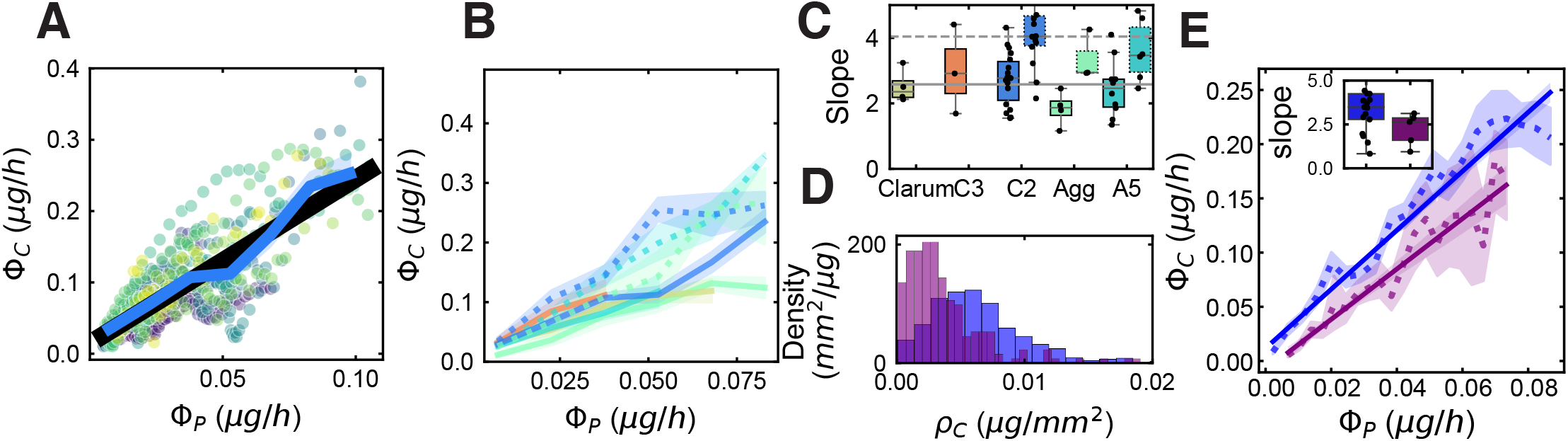
C/P exchange rate depends on plant host but not P availability. **(A), (B), (D)** Carbon expenditure rate (Φ_*C*_ ) as a function of Phosphorous supply rate (Φ_*P*_ ). **(A)** *R. irregularis* strain C2 with genotype 2 carrot root. Each point corresponds to a measurement of Φ_*C*_ and Φ_*P*_ at one timestep for one replicate, each replicate is shown in a different colour (n = 18). Blue line shade corresponds to binned average 95% C.I. over regular Φ_*P*_ intervals. Black line corresponds to linear fit over the blue points. **(B)** For *R. irregularis* A5 (cyan; *n*_genotype 1_ = 11, *n*_genotype 2_ = 6 ),C2 (blue; *n*_genotype 1_ = 18,*n*_genotype 2_ = 11), C3 (orange, n=3), *R. aggregatus* (green;*n*_genotype 1_ = 4, *n*_genotype 2_ = 3 ), *G. clarum* (yellow, *n* = 4) associated with genotype 2 (dashed line) and genotype 1 (full line). Lines and shades correspond to the same calculations as in **(A), (C)** Slopes of the linear fit forced through the origin for each host-fungus genotype combination. The box represents the interquartile range (IQR), with the central line indicating the median. The whiskers extend from the box to the minimum and maximum values within 1.5 times the IQR. Full boxes correspond to genotype 1 and dashed boxes to genotype 2. Grey line corresponds to the mean of all slopes for each genotype. Full line correspond to genotype 1 and dashed line to genotype 2. Number of replicates and color-coding is the same as in **(B). (D)** Distribution of instantaneous carbon density (*d* _carbon_) at high (1.4*µg/mL*, blue) and low (0.6*µg/mL*, purple) available phosphorous concentrations in the fungal compartment for *R. irregularis* C2 grown with genotype 2 carrot root. Each count included in the bins corresponds to a measurement of *ρ*_*C*_ at one timestep for one replicate. Number of replicates in the high- and low-P conditions were 18 and 7, respectively, totaling 722 and 209 counts, respectively. **(E)**Φ_*C*_ as a function of Φ_*P*_ for *R. irregularis* strain C2 networks grown at high (blue) and low (purple) P concentrations. Lines and points are as in **(A)**. *inset* : Slopes of the linear fit forced through the origin for each replicate for each condition. The box represents the interquartile range (IQR), with the central line indicating the median. The whiskers extend from the box to the minimum and maximum values within 1.5 times the IQR.

By contrast to this near-invariance across fungal genotypes, we found a clear change in the apparent exchange rate upon changing the genotype of the plant host (Fig. 4B,C). When grown with host genotype 2 (i.e. fast growing root), the slope of the dependence of Φ_*C*_ on Φ_*P*_ increased for all three tested fungal genotypes (*R. irregularis* A5, *R. irregularis* C2, *R. aggregatus*) compared to when grown with host genotype 1. And similarly to when grown with genotype 1, these slopes across the three fungal strains were remarkably similar, despite the stark contrasts in their growth traits (*ρ*_*C*_, *v*_wave_) when grown with either hosts (Fig. S3). Thus, while the apparent exchange rate between C and P remains constant across fungal strains when attached to the same plant host, it can vary in a manner that depends on the plant host genotype.

If a change in the plant host (which supplies C) can vary the C/P exchange rate, could varying the availability of P also have an effect? To answer this question, we repeated the experiments with different amounts of P in the fungal compartment. Our hypothesis was that, because of the reciprocal nature of exchange, lower P availability would result also in a reduced C supply from the plant, thereby rendering the C-limitation of growth more stringent. The fungus could then respond by decreasing the saturating carbon density *ρ*_*C*_, the propagation speed *v*_wave_ of its travelling-wave growth, or both.

We reduced the amount of soluble P available for fungal absorption [*P* ]_0_ to approximately one third, and found that the fungal density *ρ*_*C*_ was substantially reduced, by approximately two-fold (Fig. 4D, Fig. S3). By contrast, we found the wave speed *v*_wave_ was barely affected by this reduction in [P] (Fig. S3). This pattern of change in the travelling-wave growth phenotype implies a lower overall rate of carbon expenditure Φ_*C*_, because Φ_*C*_ is proportional to 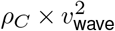 (Methods). Remarkably, despite this reduction in carbon flux, the relationship between Φ_*C*_ on Φ_*P*_ remained well-approximated as a straight line, with nearly the same slope as the control condition with 3-fold higher [*P* ]_0_ (Fig. 4E). This invariance in the C/P exchange rate motivates a parsimonious explanation for the observed reduction in *ρ*_*C*_: it reflects the response of the fungus to a reduced C budget, which in turn results from the attenuated P supply for obtaining C through reciprocal exchange.

Overall, these data are consistent with a mode of reciprocal C/P exchange in which C supply from the plant host Φ_*C*_ is approximately proportional to P supply from the fungus Φ_*P*_ . The constant of proportionality Φ_*C*_*/*Φ_*P*_ can be considered the exchange rate for reciprocal trade, and appears invariant under changes in AM fungal genotype and P available to the fungal network, but can depend on plant host genotype.

### P absorption shapes network growth phenotypes and symbiotic outcome

Can the observed proportionality of reciprocal resource exchange help us understand other salient features of AM fungal behavior, and, in turn, how these features affect symbiotic outcomes? To answer this question, we extended a recently proposed model of fungal travelling-wave range expansion (Oyarte Galvez et al., 2025) to include a dependence of network growth on reciprocal resource exchange. The model describes the coupled dynamics of growing hyphal tips and hyphal filaments at the coarse-grained level of spatial densities, in terms of microscopic parameters such as hyphal growth speed and branching rate (Methods). In extending this model to account for reciprocal exchange, we were guided by the following two hypotheses: (1) at all times, the rate at which C is received from the plant host Φ_*C*_ is proportional to the rate of fungal P transfer Φ_*P*_ ; (2) at all times, the fungal network expends all received carbon into growth.

Motivated by the observation (Fig. S3) that the wave speed remained nearly invariant under a change in P supply (which in turn limits the rate of C aquisition), we assumed the wave speed was a fixed parameter for a given plant-fungal genotype pairing. Because Φ_*C*_ increases with the rate of hyphal branching at fixed wave speed, we allowed the growing fungal network to dynamically adapt its branching rate in a manner that maintains proportionality between the rate of carbon expenditure on growth Φ_*C*_ and the rate of phosphorous transfer Φ_*P*_ . For example, when Φ_*C*_ is higher than that required to achieve the target ratio against the current value of Φ_*P*_ , branching rate is decreased. The relevant output variable of the model to implement this closed-loop feedback is the hyphal length density *ρ*, with which both Φ_*C*_ and Φ_*P*_ scale, through their proportionality to network volume and surface area, respectively.

Another important ingredient of the extended model is a dependence of hyphal width on network expansion speed. This was motivated by our finding that faster expanding strains tend to invest more in wider hyphae (Fig. S6), a relationship consistent with findings in other filamentous fungi (Chevalier et al., 2024). Because wider hyphae are more costly, this dependence implies that faster expansion comes at a higher carbon cost, and hence ought to be accounted for in a model addressing cost-benefit aspects of fungal growth strategies. We assumed the simplest affine form for the functional dependence of the average hyphal radius on wave speed, ⟨*r*⟩ = *av*_wave_ + *b*, with parameters *a* and *b* obtained by fitting the data in Fig. S6.

To test the extended model, we parameterized it with measured network-growth data and examined the predicted patterns of growth. Parameter sets for slower expanding strains (Fig. 5A, left) yielded denser growth than those for faster expanding strains (Fig. 5A, right), while the rate of phosphorus transfer (Φ_*P*_ ) and the rate of carbon expenditure (Φ_*C*_) were indeed maintained approximately proportional throughout (Fig. 5B). These results suggest that the apparent tradeoff we observed between network density and expansion speed (Fig. 2A) could be explained by travelling-wave migration under the constraint of a fixed C/P exchange rate and expansion speed-dependent hyphal thickness.

**Figure 5.**
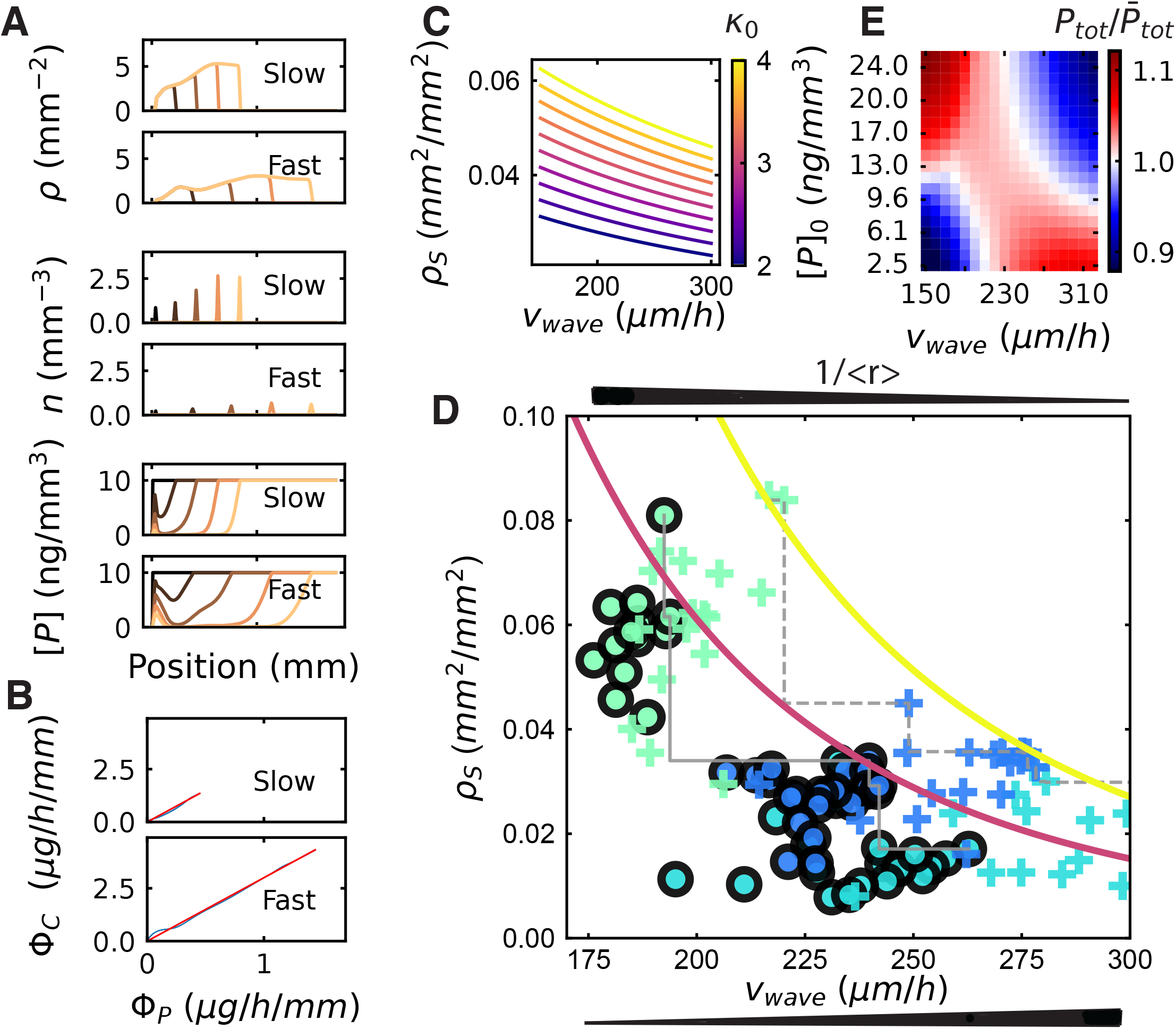
Travelling-wave model with exchange-limited growth reveals an exchange-rate-dependent Pareto front. **(A)** Model integration with *v*_*d*_ = 150*µm/h*. (upper) Model integration with for *v*_*d*_ = 300*µm/h* (lower): Colors gradients correspond to sampling at regular time interval from *t* = 0h to *t* = 900h. First row is the hyphal length density *ρ*, second row is the tip density *n* and last row is the P concentration *P* . **(B)** Φ_*C*_ as a function of Φ_*P*_ . Red line corresponds to Φ_*C*_ = *κ*_0_Φ_*P*_ , blue line corresponds to model outputs. Left and right are as in **(A). (C)** Family of “iso-exchange rate” curves computed from the analytical approximation of the travelling-wave model in the long-time limit (Methods), indicating a trade-off between surface area density *ρ*_*S*_ and wave speed *v*_wave_. These curves bound from above the range of achievable AM fungal growth strategies constrained by the value of the fixed exchange rate, *κ*_0_ (indicated by color). **(D)** Experimentally observed phenotypes in *ρS* -*v*_wave_ space. Data points represent median values over successive 10-hr intevals in each plate, with color code as in Fig. 4. Circles, to host genotype 1; crosses, host genotype 2. Gray lines, moving maximum of *ρ*_*S*_ for decreasing values of *v*_wave_ for each host genotype (full: genotype 1, dashed, genotype 2). Coloured curves, predicted Pareto fronts computed from the analytical approximation for intermediate times when the P depletion front has yet to fully develop (SI) for *κ*_0_ = 3 and *κ*_0_ = 4 (color code as in (**C**)). **(E)** 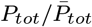as a function of wave speed *v* _wave_ and initial concentration *C*_0_. Colour corresponds to the relative increase in P absorption compared to the average of the line for a given concentration (see Methods). Integration parameters are detailed in SI.

What are the functional implications of this tradeoff? Whereas fast network expansion (high *v*_wave_) promotes exploration for new resources and trade partners, a higher network density allows for faster exploitation of local resources (because Φ_*P*_ scales with the network surface area density *ρ*_*S*_, which in turn is proportional to network length density *ρ*). The observed density-speed tradeoff can therefore be interpreted as an exploration-exploitation tradeoff: fungal phenotypes that prioritize distal exploration will tend to receive less carbon in the short term, whereas those that rapidly exploit local resources will tend to be slower at discovering new resources and trade opportunities.

Consistent with this idea, analytically solving a simplified version of the extended model (Methods) yielded a family of “iso-exchange rate” curves of negative slope in *ρ*_*S*_-*v*_wave_ space (Fig.5C, Top). They correspond to the predicted position of fungal phenotypes under the hypothesis of fixed exchange rate. These curves can be considered Pareto fronts (Shoval et al., 2012) delineating the limits of achievable travelling-wave growth strategies under the constraint of fixed exchange rate — as the C/P exchange rate *κ*_0_ is increased, the Pareto front shifts upward, thereby broadening the range of accessible strategies. Indeed, plotting our experimental data (Fig. S3) in *ρ*_*S*_-*v*_wave_ space revealed a shift in the apparent Pareto front upon swapping the plant host and therefore increasing the exchange rate (Fig.5C, Bottom). Finally, the same model could also explain the decreased fungal density at lower P availability (SI), consistent with our experimental observations (Fig. 4C). We therefore conclude that the extended travelling-wave model with exchange-rate feedback and speed-dependent hyphal widths captures salient patterns of fungal phenotype distributionss and their dependence on environmental conditions.

We further asked whether and how this exploration-exploitation trade-off could also explain environmental context-dependence of the AM symbiosis. We addressed this question by simulating fungal network growth under different combinations of initial P concentration [*P* ]_0_ and fungal growth strategy *v*_wave_ . For each initial concentration, we compared the total amount of P transferred during a fixed interval of time *P*_*tot*_ to its average 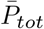 over all strategies. The resulting ratio is a measure of P absorption performance and hence reflects also the comparative benefits of different fungal strategies for P supply to the plant. We found that the wave speed that provides the highest relative phosphorous transfer 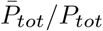 depended on [*P* ]_0_ (Fig. 5D). Specifically, at higher [*P* ]_0_, slow-expanding networks that invest in dense growth without paying extra costs can better capture the available P. By contrast at low [*P* ]_0_, fast-expanding networks can better escape their self-generated P-depletion zone (Fig. 5A, bottom panels), leading to an overall advantage in capturing P.

These results demonstrate how the P absorption and transfer performance of AM fungi, which often determines observable symbiotic outcomes such as plant growth (Corrêa et al., 2024), can strongly depend on the context of symbiosis, such as the plant-fungal genotype combination (which strongly affects *v*_wave_) and environmental conditions such as the accessible P concentration.

## Dicussion

By leveraging robotic imaging combined with machine learning, we developed a new approach to estimating resource fluxes in the mycorrhizal symbiosis based on *in toto* quantification of network morphology throughout growth. We discovered a systematic proportionality between carbon transfer from the plant host and phosphorus transfer by the AM fungal network (Fig. 4). We found that the proportionality coefficient remained constant across various fungal strains associated with the same plant host, despite wide variation in their growth phenotypes. Collectively, our results suggest that fungal network growth is constrained not by carbon supply per se, but rather the exchange rate of reciprocal C/P exchange, which determines the amount of carbon that the fungus can obtain per unit of collected phosphorus.

Previous studies that used isotope tracers in similar *in vitro* root organ cultures (Kiers et al., 2011) and in real soils (Lekberg et al., 2024) obtained compelling evidence for reciprocal exchange, but whether and to what extent the exchange is proportional has remained challenging to determine. In *in-vitro* cultures, it was found that more host-derived carbon was transferred to fungal compartments where a greater quantity of P was available (Kiers et al., 2011). In real soils, quantitative radiotracer counting showed a positive correlation between P delivery to the plant and host C allocation to the AM fungal partner (Lekberg et al., 2024). However, it has also been suggested that P supply may not always be tied directly to carbon allocation. Specifically, C surplus theories hypothesize that the amount of carbon transferred to mycorrhizal fungi is independent of nutrient return from the fungus (Bunn et al., 2024) and only depends on plant metabolism and fungal “carbon sink strength”. Our present work demonstrates that fungal “carbon sink strength” is not fixed by the size of the network. Instead, both range-expansion speed and saturating network density affect how much carbon the fungal network consumes at a given point in time (Fig. 2, Fig S3). We also found a trade-off between network density and expansion speed. This is important because it suggests that the fungal partner is constantly constrained by carbon exchange, such that the same network cannot be both dense and fast.

Our finding that the C/P exchange rate depends on the host plant genotype raises a wealth of interesting questions. How might this exchange rate vary with various environmental conditions of the host? And might the rate vary if the plant were trading with not one, but multiple fungal partners simultaneously? More generally, what are the factors that set the exchange rate, and through what mechanism do they act? On the host side, control of arbuscule formation (Paries and Gutjahr, 2023) or lifetime (Javot et al., 2007; Gutjahr and Parniske, 2017) have been suggested as potential mechanisms to control carbon flux into AM fungal networks. The necessity for control of lipid biosynthesis and export to the AM partner may also require complex feedbacks involving regulation of gene networks (Zhang et al., 2023).

The context dependency of the plant-mycorrhizal symbiosis has been the focus of research for decades (Pearson and Jakobsen, 1993; Smith et al., 2003; Lendenmann et al., 2011; Koch et al., 2017; Lee and Eom, 2015; Hoeksema and Schwartz, 2003; Hoeksema et al., 2010). Despite this growing body of work, predicting the nutrient-exchange outcomes of the symbiosis has remained a puzzle. Our experimental findings and theoretical model provide a new foundation for exploring how proportionality of resource exchange affects total symbiotic outcomes, such as P transfer from AM fungi to plants, across varying contexts and hosts.

Our modelling framework could also help understand how changes in the mobility of nutrients such as P, due *e*.*g*. to layered soil texture or chemical properties, can affect the optimal range-expansion strategy. Typically, root systems are simultaneously colonized by multiple AM fungal partners (Zhou et al., 2020; Vandenkoornhuyse et al., 2002). It would also be straightforward to extend our modeling framework to include multiple species with contrasting range-expansion strategies, to address the stability of coexistence (Gude et al., 2020). An important question is how a diversity of partners with complementary traits could affect P transfer to plants, and whether the proportionality documented here changes as the number of partners change (Wyatt et al., 2014). Exploring these questions could help explain why so many AM fungal species can co-exist within the same niche, a long-standing question in AM ecology (Camenzind et al., 2024).

Future work is needed to simultaneously characterize these traits across fungal strains found in different environments, as well as quantifying the total amount of P transferred to plants. For example, in previous pot experiments grown under different P fertilization treatments, *R. irregularis* provided more P to the host plant compared to *R. aggregatus*. However, the difference between the two strains became insignificant with higher P fertilization (Wang et al., 2016). Following the temporal dynamics of these differences, and how they coincide with amount, location and regularity of fertilizer applications is a next important step.

Our work demonstrates the functional significance of subtle, micron-scale traits, such as the hyphal radius distribution, in understanding symbiotic resource exchange. We initially suspected that thinner structures would be favoured for resource absorption. This is because the carbon cost of a hypha grows as the square of its radius, while its ability to absorb nutrients from the environment grows only linearly. The most cost-efficient absorbing structures therefore are the thinnest ones. This principle is exemplified in plants where root hairs are thinner than main roots, and also in AM fungi where branched absorbing structures (BAS) (Bago et al., 1998) tend to be the thinnest of all hyphae across the AM network (Fig. 1C). However, our findings on how hyphal radius relates to other growth phenotypes indicates that minimizing the cost of building absorption structures is not the only optimization objective. We found that faster expanding networks tend to invest more in thicker hyphae (Fig. S6). This suggests that fast range expansion requires investment in thicker runner hyphae to allow the transport of C to the network periphery, as well as the transfer of P back to the host root. Our observation that AM fungi can change their hyphal width over time (Fig. 1E) further raises the possibility that hyphal width is a dynamically regulated morphological trait - perhaps to enhance the efficiency of long-distance nutrient transport necessitated by their symbiotic lifestyle.

Because our data were gathered under sterile and *in-vitro* laboratory conditions, it remains unknown whether, and to what extent, the observed proportionality of reciprocal exchange will extend to symbioses under natural conditions in which hyphae have their own microbiome that contribute to nutrient uptake (Vaishnav et al., 2025; Berrios, 2025). It is also well known that AM fungi provide plants with a diversity of other benefits, including water, nitrogen, protection from pathogens, etc. (Martin and van der Heijden, 2024). Based on our data, we would anticipate that benefits (e.g. nitrogen) that rely on network morphology, such as the total absorption surface area, might also yield similar proportionalities in the pattern of exchange. By using a similar approach to our phosphorous measurements, other nutrient exchanges of the symbiosis could be tested. Density and wave speed can also be important traits for the protection against pathogens because of competition for nutrients (Lutz et al., 2023). Dense mycelium could effectively create a nutrient depletion barrier around the root that would make the host less accessible to pathogens.

In summary, our results identify the carbon-phosphorus exchange rate as a key constraint on AM fungal growth and range-expansion strategies. We found AM fungal genotypes were not positioned arbitrarily in the space of all possible strategies, but within a subregion bounded by an exchange-rate-dependent Pareto front (Fig. 5D). The shapes of the observed Pareto fronts suggest that fungal range-expansion strategies trade off exploration and exploitation performance under a given exchange rate. In turn, how fungal strategies position themselves within this phenotype space impacts P-transfer outcomes to the host plant. These observations provide a foundation for a mechanistic understanding of the stability of the mycorrhizal symbiosis, and also for predicting coupled P and C dynamics at the earth system scale. Finally, the mass exchange ratio of ≈ 3 C/P implies that the flux of carbon in one direction and that of phosphorus in the other are of a similar order of magnitude. Maintaining mass fluxes of comparable magnitudes in opposite directions throughout the AM fungal network must require sophisticated strategies for routing resources throughout the network and perhaps even at the level of individual hyphae. How these dedicated symbionts manage their ‘supply-chain dynamics’ for reciprocal nutrient exchange is a promising direction for future investigations.

## Methods

### Sample preparation

### General recipe

We used modified Strullu-Romand (MSR) medium(Declerck et al., 2005) solidified with (3g/L) Phytagel and containing 10g/L sucrose (Carl Roth). The MSR medium in this study contained final concentrations of 739 mg/L MgSO_4_.7H_2_O, 76 mg/L KNO_3_, 65 mg/L KCl, 4.1 mg/L KH_2_PO_4_, 359 mg/L Ca(NO_3_)_2_.4H_2_O, 0.9 mg calcium pantothenate, 1 *µ*g/L biotin, 1 mg/L nicotinic acid, 0.9 mg/L pyridoxine, 0.4 mg/L cyanocobalamin, 3 mg/L glycine, 50 mg/L myo-inositol, 1.6 mg/L NaFeEDTA, 2.45 mg/L MnSO_4_.4H_2_O, 0.28 mg/L ZnSO_4_.7H_2_O, 1.85 mg/L H_3_BO_3_, 0.22 mg/L CuSO_4_.4.5H_2_O, 2.4 *µ*g/L Na_2_MoO_4_.2H_2_O, and 34 *µ*g/L (NH_4_)Mo_7_O_24_.4H_2_O.

In the root compartment, the phosphate content of the medium was reduced to 1% of the above-mentioned concentration (41 *µ*g/L KH_2_PO_4_) to stimulate mycorrhizal colonization of the roots. In the fungal compartment, we adjusted the phosphorus content in the MSR medium to impose low-P and high-P conditions:we applied either a high-P condition using the full MSR medium as described above, or a low-P condition treat in which KH_2_PO_4_ was removed entirely.

### Low- and high-P conditions

Specifically, for the low-P condition, we excluded KH_2_PO_4_ from the medium. Phytagel, the gelling agent used in this study, however, also contains P in a concentration that could not be modified. In preliminary experiments using Spectrophotometric assays (see SI), we estimated the concentration of P in the phytagel to be 27 *µ*mol/g. The fungal compartment was composed of 28mL of MSR medium. Based on the above concentrations, this means the fungal compartment contained 96 *µ*g P in the high-P condition and 70 *µ*g P in the low-P condition corresponding respectively to respective concentrations of 3.4 *µ*g/mL and 2.5 *µ*g/mL.

We observed that the concentration of P in the root compartment did not seem to significantly decrease over time (Fig. 3b). Also, observations over longer timescales ( ≥ 10 days) showed that the concentration in the fungal compartment seemed to plateau around similar but slightly higher value as the root P concentration after its initial decrease. P is known to be bound/adsorbed by soluble aluminum, iron, and manganese at low pH(Etesami et al., 2021). pH in our plate was around 5, suggesting that potentially a large proportion of the P we could measure was not available to the plant or fungus. The slightly lower plateau concentration in the root compartment could be due to a change in pH due to the root presence. Based on measurement of root compartment plateau P concentration and fungal compartment plateau concentration at time ≥ 10 days, we estimated that 2 *µ*g/mL of P were inaccessible. 0 concentration of P therefore corresponds to 0 concentration accessible by fungus and root but 2 *µ*g/mL of P measured with our method. Concentration accessible by the fungus are therefore 1.4 *µ*g/mL in the high-P condition and 0.5 *µ*g/mL in the low-P condition.

#### Fungal Strain

All experiments were performed with *Rhizophaghus irregularis* strains A5 DAOM664344, C2: DAOM664346, C3 (Ian Sanders’ lab, Lausanne, Switzerland), *Rhizophaghus aggregatus* and *G. clarum* (Ian Sanders’ lab, Lausanne, Switzerland). All AM fungi were cultivated on regular MSR medium associated with Ri T-DNA transformed carrot root for 2-6 months until plates were fully colonized.

#### Root Strain

All experiments were performed with two different root organ culture of Ri T-DNA transformed carrot root. Ri T-DNA transformed carrot root “EU” (referred to as as genotype 1 in main text) originated from a lab in Europe (see Oyarte Galvez et al. (2025)). Ri T-DNA transformed carrot root “CA” (referred to as as genotype 2 in main text) was Daucus carota subsp. Sativus, Cultivar enterprise and originated from Canadian Collection of Arbuscular Mycorrhizal Fungi and was derived from Clone P68.

### Estimation of hyphal radius from low resolution images

#### Label dataset generation

Fungal networks grown in control conditions were imaged in the timelapse imaging system every 2 hours (see SI and (Oyarte Galvez et al., 2025) for details). Immediately following imaging, we transferred the plates to the high-resolution imaging microscope. We then imaged locations across the network and recorded their network position in in high resolution (i.e. 50X, see SI). We focused by positioning the optical plane in the centre of the hypha. This was done on the basis that the cell wall should look thin and the active flows in the interior should be the widest. For each plate, we were able to accomplish this in less than 2 hours. Images were then labelled using the software labelme. We drew three lines across the hypha that would span its whole external diameter. This external diameter included the hyphal cell wall. Using past experience, it was possible to establish what part of the imaged hypha corresponded to the interior with active flows and that attributed to the dark thick cell wall. In general, the dark thick cell wall never contributed more than 10-20% of the total diameter, which is in line with previous estimates of hyphal cell wall thickness for AMF(Bago et al., 1998). For each hyphal segment, we assigned the diameter corresponding to the mean of the three measurements. By repeating several independent measurements along several edges on these high-resolution image, we estimated the standard deviation of the manual estimate to be of 0.3*µ*m. This variation was either due to the difference in appreciation of the exact boundaries of the hyphae on the high-resolution image or to effective biological variation of the hyphal radius along an edge in a single field of view.

#### Feature dataset generation

Next we segmented and skeletonized the networks as in (Oyarte Galvez et al., 2025). For each edge that had been imaged in higher magnification and labelled, we manually identified the corresponding edge of the network. We generated transects perpendicular to the skeleton and of size 120 pixels with the function profile_line of the package skimage (Van Der Walt et al., 2014). We selected the transect that was the closest to the position of the labelled image to use as a feature.

#### Data augmentation

For part of the dataset, we varied the focus over 0.1 to 0.2mm above and below the “optimal focus” for images and varied illumination. Each label from high magnification was therefore associated with multiple features corresponding to the different focus. In some other datasets, we extracted multiple transects close to the high magnification labelled image and associated them with the same label. This data augmentation procedure was done to help the model generalize over different focus and illumination settings. We also added transect corresponding to regions of the image empty of any hypha and a zero radius corresponding values in order to limit the overestimation of thinner hyphae.

#### Final dataset

The final dataset, following data augmentation consisted of 3182 elements, which were divided into training, validation, and testing sets. Specifically, 90% of the data (approximately 2,546 samples) was used for training and validation, while the remaining 10% was kept for final testing, resulting in 292 samples for the test set. The test set consisted in a combination from entirely independent timesteps and plates from the rest of the dataset. This means, none of the data corresponding to the plate at that timestep was used during training and completely new plates were included in this test set. We resampled the test set so it would follow the same distribution as the training set and comparison of errors would be comparable. To do so, we calculated the frequency distribution over 20 bins of the training set and sampled with replacement from the full test set. The distributions of the training/validation and resampled test sets are shown in Fig. S1 A,C. Values ranged from 0 to 8 in both cases.

### Carbon cost and phosphorous absorption calculations from data

#### Total Network Carbon

For a given edge *i*, of length *L*_*i*_ as measured by segmentation, and radius *r*_*i*_ as estimated with the CNN model, we approximated each edge to be a cylinder and therefore its volume *V*_*i*_ to be 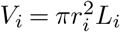. The total mass of carbon in the entire hyphal network at a given time *m*_*C*_(*t*) can then be estimated as:

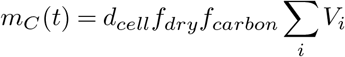

where *d*_*cell*_ is the mass density of the hyphal cell, *f*_*dry*_ is the dry mass fraction of the total (wet) mass, *f*_*carbon*_ is the fraction of dry mass accounted for by carbon, and the sum runs over all edges *i* of the network at time *t*. We chose *d*_*cell*_ = 1.1 *g/cm*^3^ (typical value for cells (Bakken and Olsen, 1983)), *f*_*dry*_ = 21% (typical value for fungi (Bakken and Olsen, 1983)), *f*_*carbon*_ = 50% (typical value for cells (Bar-On et al., 2018)).

We followed a similar principle to compute the mass of carbon in each spore. We first computed the volume of each spore from its radius estimated with classical image analysis techniques (see Oyarte Galvez et al. (2025)). In lieu of known data for spore-specific values for *d*_*cell*_, *f*_*dry*_, and *f*_*carbon*_, we assumed the same values as those for hyphae. Total network carbon 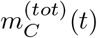 was computed by summing the volume of all individual edges composing the network and all the spores at each point in time.

We note that the carbon-estimation parameters *d*_*cell*_, *f*_*dry*_, and *f*_*carbon*_ used here should be considered estimates for a generic fungal cell, and hence do not reflect exact values for AM fungi. Yet in lieu of definite evidence for or against these values within the AM fungal literature, we take these generic estimates as our current best approximation. Indeed, previous study also used very similar estimates for the same parameters for AM fungi (Jakobsen and Rosendahl, 1990; Bar-On et al., 2018). We also note that these parameters can in principle vary across hyphae within a given network and over time. Yet given this formula for *m*_*C*_(*t*) sums over all hyphae of the network, these parameters can be considered network-wide averages that should be quite stable in time during steady travelling-wave growth (Rautio et al., 2006; Saldanha et al., 2004). The limitations of these simplifying assumptions are discussed in further detail in SI.

#### Carbon Density

To compute the carbon density, we calculated the change in area Δ*A* and in total network carbon 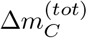 over a time interval Δ*t*. The carbon density was then defined as:

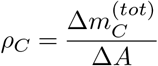

In the case of Fig. 2A we choose Δ*t* = 10*h*.

#### Rate of Carbon Expenditure

Rate of Carbon Expenditure was defined as:

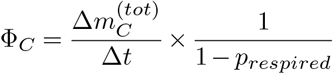

where *p*_*respired*_ is the proportion of carbon that is respired during growth. It is equal to 1 − *CUE*, where *CUE* is carbon use efficiency i.e. the proportion of carbon incorporated in biomass and not respired of all the carbon used, a quantity often measured in ecology. We used an average CUE for soil microorganisms of 50% (Manzoni et al., 2012). It is important to note that CUE reported in the literature can encompass variable definitions and measurement methods. We didn’t find any specific estimates for Arbuscular Mycorrhizal Fungi that would match our criteria. In nutrient poor Boreal forests, CUE of ectomycorrhizal fungi (EMF) was found to vary between 3 and 15% (Hagenbo et al., 2019). We however expect AMF CUE to be significantly higher since, on the contrary to EMF, they invest no carbon in substrate decomposition. While our estimate of CUE for AMF is uncertain, it does not affect the main conclusion of the paper since a different CUE, if it is not strain specific, will not break the observed proportionality between P absorption by AMF and C use but simply change the proportionality factor.

#### Computing P flux into the network

**P flux in the low P regime** Computing Φ_*P*_ in the low P regime necessitated evaluating Eq.3 at every time point during network growth. To do so, we implemented a three-dimensional explicit finite-difference numerical integration scheme to simulate reaction-diffusion dynamics of phosphorus (P) within a semi-circular petri dish geometry. The computational domain was discretized into a structured grid with dimensions of 30 × 30 × 5 grid points in the x, y, and z directions, respectively, spanning horizontally and to a depth of 8.8 mm, representing the agar layer. The top layer of cells (i.e. z=0) had a non-zero reaction term that followed Michaelis–Menten kinetics:

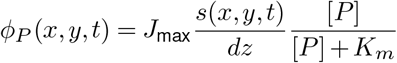

where *s*(*x, y, t*) represents the local fungal surface area density (surface area per unit area of agar surface). Values of *s*(*x, y, t*) were derived from experimental measurements of fungal surface area across 18 distinct regions of the petri dish, each divided by the area of their respective regions.

The final equation that is being integrated is therefore:

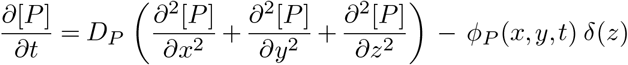

Where Dirac delta *δ*(*z*) represents the fact that absorption only happens at the surface.

Initial phosphorus concentration was uniformly set to 0.5*µgP*.*mm*^−3^ across the domain. Diffusion was modelled isotropically with a diffusion coefficient *D*_*P*_ = 1.8*mm*^2^.*h*^−1^ corresponding to the one estimated for phosphate in Agar gels at 25°C (Davison, 2016).

Zero-flux (Neumann) boundary conditions were enforced laterally at the geometric boundaries of the semi-circular dish and vertically at both the top (surface) and bottom of the agar layer.

Each numerical integration was initiated at a virtual time *t*_0_ before the start of imaging where the total area of the network was estimated to be 0. *t*_0_ was estimated by fitting an affine function to the first 5 measured values of area for each plate and finding the time at which the affine function crossed the the y= 0 line. We then extrapolated linearly the measured surface area in each region up until *t*_0_. Such extrapolation was bounded to ensure non-negative values.

Experimental surface area measurements were interpolated linearly in time, decoupling the numerical integration timestep from the experimental imaging frequency.

Numerical stability was ensured by selecting the time step sufficiently small compared to the spatial meshing

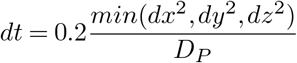

Concentration fields were updated iteratively using an explicit Euler scheme, integrating from *t*_0_ to the end of the experiment, capturing transient concentration dynamics and total phosphorus flux. Code for this integration is available in the following repository together with the rest of the replication package.

The total phosphorus flux was computed by integrating reaction terms over all cells.

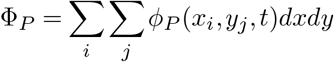

#### P flux in the high P regime

In the regime where [*P* ] is high enough that P transport across the membrane works at saturation (i.e. [*P* ] ≫ *K*_*m*_), P depletion proceeds proportionally with the time-integrated surface area 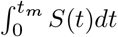 *dt* as in Eq. 2 and computing the P flux Φ_*P*_ simplifies from Eq. 3 to 1. This is justified by the fact that all data points correspond to values of the integrated surface area that are well below 10^10^*µm*^2^*h* and therefore correspond to a regime where this equation is valid according to Fig. 3C. All values 4A,B were therefore computed using Eq. 1.

### Model of network propagation with feedback

#### General Framework

The general framework is the same as explained in (Oyarte Galvez et al., 2025), the growing tips are represented in terms of their spatial density *n*(*R, t*) (with units of number per 3-dimensional volume, e.g. [mm^−3^]). To represent the hyphal filaments laid down by growing tips, we introduce an additional variable *ρ*(*R, t*) to represent their spatial density (with units of filament length per 3-dimensional volume, e.g. [*µ*m/mm^3^]).

To represent the fact that tip growth lays down hyphal length at a rate *v*_g_ corresponding to the average speed of hyphae that densify space we set

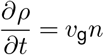

Then we represent the local creation and annihilation of tips

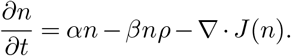

Where *α* is the branching rate, *β* the anastomosis rate and *J*(*n*) is a spatial flux that represents the movement of tips across the unit volume (see SI).

#### Accounting for radius differences

In order to convert a total length of fungal hyphae into the corresponding network volume we use the length weighted mean radius ⟨*r*⟩ or root mean square ⟨*r*⟩_*RMS*_ where

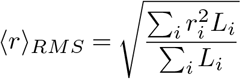

which can be rewritten

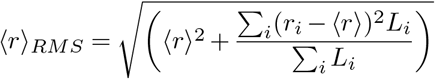

Setting 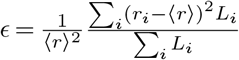 which corresponds to the square of the coefficient of variation of the radius distribution we obtain.

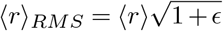

At first order, *ϵ* does not strongly depend on the genotype (see Fig. 1F) and we can therefore set *ξ* = 1 + *ϵ* which is considered to be a constant *ξ* ≈ 1.6.

We observed that hyphal extension speed is constrained by radius (Fig. S6A), a relationship consistent with findings in other filamentous fungi (Chevalier et al., 2024). Averaged over whole networks, this translated into slower growing networks having smaller ⟨*r*⟩ (Fig. S6B) and ⟨*r*⟩ _*RMS*_ (Fig. S6C).

This suggests that the hyphal radius ⟨*r*⟩ used for model integration should also vary with speed.

Because even more than five strains are needed to precisely estimate the functional form to this dependence, we can instead give a generalist functional form ⟨*r*⟩ = *av*_wave_ + *b* with *a* = 5.0 × 10^−3^*h* and *b* = 1.3*µm* found by fitting an affine function to the minimum ⟨*r*⟩ as shown in Fig. S6B.

#### Carbon Cost and Network P Absorption from mode

By integrating over space and assuming cylindrical symmetry around the root, we can define the network P absorption flux per unit length as:

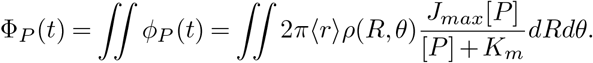

At any point in time and space, the amount of carbon used for network growth is proportional to the amount of newly built network volume. Newly built network volume in a region of space in radial coordinate is proportional to 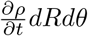. The proportionality factor *γ*_*C*_ between this newly built network volume and the amount of carbon used is the product of four parameters: mass density of the cell *d*_*cell*_, the fraction wet mass accounted for by dry mass *f*_*dry*_, the fraction of dry mass accounted for by carbon *f*_*Carbon*_, and the inverse of carbon use efficiency 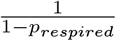 so that

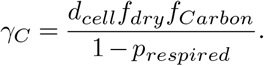

By integrating over space we can define the network carbon building cost per unit root length Φ_*C*_ as

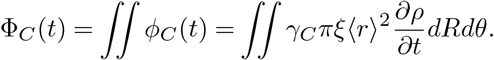

#### Exchange rate driven adaptation

The ratio 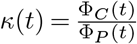 is the instantaneous exchange rate of Phosphorous in carbon units. We assume that the network adapts its growth parameter to match an average objective exchange rate *π*_0_.

Specifically, we assumed that the branching rate *α* is changing over time in order to reach the objective. This is motivated by the fact that branching in fungi is thought to be the consequence of vesicle accumulation at the tips beyond a maximum threshold. In other terms, when the supply of vesicles exceeds their capacity to be incorporated into the existing tip, they accumulate leading to the formation of a new tip (Harris, 2008). Assuming carbon availability is the limiting factor for vesicle production, it is therefore possible that AM adapt to shortage or abundance of carbon by adapting their branching rate. Since the saturation density is a growing function of *α*, at a fixed propagation speed, increasing Φ_*C*_ tends to increase together with *α*. We therefore assumed that, when the average recent exchange rate 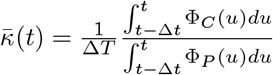 overshoots *κ*_0_ , *α* is reduced. Such an integral formulation was chosen to damp oscillatory behavior due to adaption of branching rate overshooting the objective exchange rate and we chose Δ*t* = 10*h*.

We added one last closing equation to the system:

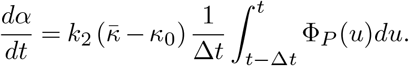

With *k*_2_ a rate parameter. Such a form for adaptation of *α* is entirely subject to a number of hypotheses. The main conclusions are however not affected by the specific form of the adaptation as long as it allows the exchange rate to match the set exchange rate on average (Fig. 5B).

***Fig. 5D***. For each concentration and each wave-speed we computed the total P transferred to the host plant *M*_*P*_ by integrating Φ_*P*_ over time: 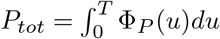. Then at a fixed initial P concentration [*P* ]_0_, we computed 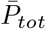, the average total transfer of P across all wave-speed sampled. The relative increase in P absorption was 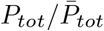

#### Analytical considerations on the proportionality between Φ_*C*_ and Φ_*P*_

For a netork growing as a travelling-wave (entirely determined by its saturation density *ρ*_L_ and its wavespeed *v*_wave_), proportionality between Φ_*C*_ and Φ_*P*_ is expected to happen in the permanent regime where a P depletion front has developed. In that regime, for a network of size *R* in radial coordinates, only a ring of size Δ*R* is exposed to regions that are not yet depleted in phosphorus. The area of that region is approximately *πR*Δ*R* and its rate of P absorption is approximately

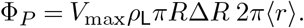

Δ*R* is directly linked to network propagation speed and can be estimated to be 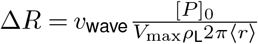 yielding

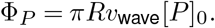

The rate of carbon investment, on the other hand, can be easily estimated to be

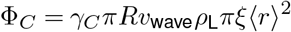

And

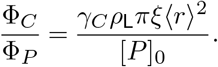

The above ratio is indeed time independent (as long as the radius *r* is constant in time). In the case where the ratio 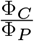 is imposed by the plant to be equal to *κ*_0_ we obtain

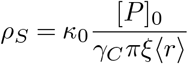

Where *ρ*_*S*_ is the surface area saturation density. Since ⟨*r*⟩ is an increasing function of *v*_wave_, this imposes immediately a trade-off between *ρ*_*S*_ and *v*_wave_. Such trade-off is represented in Fig. 5C. Although our data does not correspond to this permanent regime, the above derivation helps understanding how the trade-off can emerge and the dependency of saturation density on initial phosphorous concentration. In order to understand the variation as it is observed in our experiment (Fig. 5C, lower), we use a different framework (see SI).

### Analytical considerations on the optimal wave speed

The results shown in Fig. 5D can be understood from an analytical perspective.

#### Low P, Permanent regime

In the permanent regime, letting *R* = *v*_wave_*t* within the expression for Φ_*P*_ derived above, we have

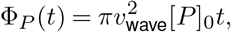

which means the fastest expanding strains (with high *v*_wave_) will always provide more *P* . This is understandable since the permanent regime corresponds to a regime where escaping the *P* depletion zone is essential.

#### High P, Exponential regime

At high P ([*P* ] *>> K*_*m*_), for a growing network with fixed average hyphal radius ⟨*r*⟩ , equating the carbon expenditure on network growth Φ_*C*_ with the phosphorus flux Φ_*P*_ scaled by a fixed exchange rate *κ* yields

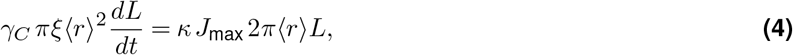

which is a differential equation describing dynamics of the total network length *L*. This equation has the solution

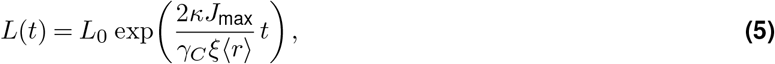

where *L*_0_ is the network length at time *t* = 0. If we choose *L*_0_ so networks of different average hyphal radii always start with an equal volume *V*_0_ at *t* = 0 we have

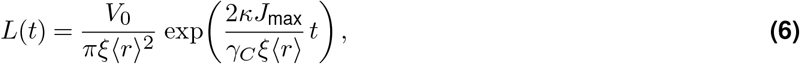

Integrated over time we obtain

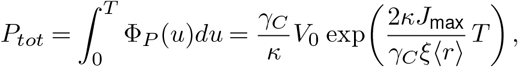

Which is a decreasing function of ⟨*r*⟩ meaning that strains with smaller radii (i.e. slower wave speed) will always achieve higher P-uptake performance in this regime.

## Aknowledgements

We acknowledge HFSP RGP (0029) to E.T.K. and T.S.S.; ERC-Nuclear Mix (101076062) to V.K.; ERC grant 834164 to S.W.; and the Grantham Environmental Trust, Schmidt Family Foundation, Paul G Allen Family Foundation, Ammodo Foundation, Hefner Foundation, Quadrature Climate Foundation, Bezos Earth Fund, NWO-VICI (202.012), NWO-Spinoza (SPI.2023.2) and NWO-MICROP (024.004.014). We thank I. Sanders for sharing fungal strains.

