## Supporting information for "Carbon-phosphorous exchange rate constrains growth of arbuscular mycorrhizal fungal networks"

#### Supplementary methods

##### Plate preparation details

***Assembly of Mesh Frames.*** Frames were designed slightly trapezoid to fit next to the central barrier of two-compartment split plates (Greiner Bio-One) with a longer top edge (88 mm) than bottom edge (85.5 mm), and a consistent height (12 mm). They include a central opening (50 x 2 mm) distanced 2 mm from the top edge. This opening connects to the upper edge of the central barrier of the plate, therefore extending the central barrier at the outsides of the plates. The fungus could cross through the opening into the second compartment. To keep the root from crossing over, the opening was covered with nylon mesh.

Acrylic frames were cut using a laser cutter. Nylon mesh (pore size 50 $\mu$m, 9 x 71 mm) was attached to the acrylic frame using UV resin in such a manner that the frame opening was fully covered by the mesh and free of resin. The resin was cured with UV light for 3 min, the frames then wrapped in aluminum foil and sterilized at 80°C for 72 hours.

***Two-compartment Split Plate Preparation.*** In a laminar airflow hood, one compartment of a sterile two-compartment split plate (94 mm diameter, Greiner Bio-One) was filled with 28 ml MSR medium. An autoclaved sheet of cellophane (Hoefer™ TE73, semi-circle with trapezoidal overhang at the straight edge) was placed on top of the solidified medium and the cellophane overhang was folded into the empty second compartment. A custom acrylic frame was inserted into the empty compartment, securing the cellophane overhang between the acrylic frame, central barrier, and the bottom of the plate. To avoid dislocation of cellophane and/or frame, 5 ml 1%P MSR was poured into the second compartment to immobilize the components. The compartment was then filled to a total of 25 ml 1%P MSR.

***Inoculation.*** In a laminar airflow hood, 2-3 cm of in vitro Ri T-DNA transformed root were transferred to the split plate compartment not covered by cellophane ('root compartment'). A circular plug containing only mycelium and spores was cut from the AMF stock plate (2-6 months). Roots were carefully removed if necessary. The inoculation plug was placed on top of the root, covering not more than half of the root. The plates were sealed with parafilm and stored horizontally and u pright in an incubator at 25°C.

***Plate Maintenance.*** Plates were controlled regularly for fungal and root growth. Any root crossing from the root compartment into the cellophane-cover fungal compartment were pulled back or removed. It usually took 2-4 weeks for the fungal network to start crossing to the fungal compartment and it was kept in the imaging system for 3-6 weeks.

#### Phosphorus Measurements

Measurements and data points are the same as the ones shown in (Oyarte Galvez et al., 2025). Details are given below.

***Agar and root preparation.*** We calculated phosphorus concentrations in fungal and root compartments by first removing cellophane from the fungal agar, and then cutting the agar into equal sized pieces (up to 18 pieces). Each agar cube corresponded to a spatial position either away or close to the root. We cut the root compartment agar into two pieces and carefully separated the root from the root agar. We weighed each agar piece and placed it in a Teflon cylinder, and we placed each root in a kraft envelope. We put the Teflon cylinders and envelopes in the oven at 70 ℃ for two days to dry. After drying, we weighed roots and placed them in a Teflon cylinder.

***Digestion.*** We added 0.5 ml of digestion mixture (HNO3/HCl 4:1) to the Teflon cylinders using a repeating pipette and left the cylinders open for 30 mins to release gases. We then placed the closed cylinders in a destruction oven at 140 ℃ with the temperature limit set to 160 ℃ for 7 hours. We opened the cylinders, added 2 ml of demiwater using a dispenser and transferred the contents of the cylinder to a test tube. We left the test tubes in a fume cupboard for at least one day to release acid fumes and covered them with plastic foil before placing them in the cold room for a week.

***Spectrophotometric determination of phosphate content.*** The phosphate estimation was based on the formation and reduction of phosphomolybdate. Following the method of(Murphy and Riley, 1962), we pipetted 150$\mu$L of the solution obtained after digestion in test tubes and added 4mL of colour reagent. The colour reagent was prepared in a 1L water solution with 13.33mL concentrated H2SO4, 1.14g ammonium heptamolybdate, 1.00g ascorbic acid and 0.026g of potassium antimony. The test tubes were left for 30 mins for the colour to form. We measured absorbance at 880nm in a spectrophotometer using plastic cuvettes. Because P is known to be bound/adsorbed by soluble aluminium, iron, and manganese at low pH(Etesami et al., 2021), we calculated that $\sim$2$\mu$g/mL of P was inaccessible in root and fungal compartments, which we used as our baseline when using equation 3.

#### Timelapse imaging and network segmentation

Imaging and general network segmentation was as described in (Oyarte Galvez et al., 2025). The rate of false negative was higher for very thin edges especially when focus conditions are not ideal. Such an effect was however estimated to be of small magnitude for most strains. In the case of *R. aggregatus*, a higher proportion of very thin edges were not detected. We therefore developed a machine-learning based approach for segmenting the plates from this strain. Such an approach yielded similar results on other strains (SI Fig. 7) but was better at detecting thinner edges (SI Fig. 8. Details of the method are given below.

***Label and feature dataset generation.*** Groundtruth images used for training were obtained from the classic (i.e. non-machine learning) segmentation described in (Oyarte Galvez et al., 2025), and manually checked by eye for quality. For a set of n=2309 images of *R. irregularis* C2, a segmentation mask (i.e. a binary image where pixels were 1 for hypha, 0 for background) was computed. The raw image was used as input and the segmentation mask as the target output. The full dataset was split into a training set of 1797 images, and a test set of 512 images. In order to mimic typical challenges posed by real data, we employed data augmentation where a random 50% of the training images were blurred, and a random 50% had intrusive background contrast patterns added to them. Although intrusive background contrast patterns were not a major issue for the data considered in this study, they were included in order to train a general purpose AMF segmentation algorithm that is robust to experimental conditions that include such background irregularities.

***Network architecture and training procedure.*** We used the U-Net CNN architecture (Ronneberger et al., 2015) as it has achieved impressive results in segmenting biological networks of similar morphology to our fungi (IV et al., 2025). U-Net alternates convolutional filters with both max-pool layers, where the image is downsampled by taking only the maximum pixel value in a $N \times N$ region (in our case $N = 2$), and dropout layers. Developed as a method to avoid overfitting, dropout layers temporarily remove a random subset of connections from the network at each training step (Srivastava et al., 2014). As such, U-Net first compresses, and then expands the data, resulting in a U shaped architecture. The network also contained skip connections with the aim of preserving large-scale features from the contracting path in the expanding path. Our full model consisted of 31,036,480 trainable parameters.

The network was trained using Stochastic Gradient Descent with Nesterov Momentum (Sutskever et al., 2013) according to

$$\theta_{t+1} = \theta_t + v_{t+1}, \quad (1)$$

$$v_{t+1} = \mu v_t - \eta \nabla \mathcal{L}(\theta_t + \mu v_t), \quad (2)$$

where $\theta_t$ are the network parameters at timestep $t$, $\mu = 0.99$ is the momentum coefficient, $\eta = 10^{-4}$ the learning rate, and $\mathcal{L}$ the loss function calculated over a batch. We used a batch size of 1. By a process of trial-and-error experimentation we chose a loss combining the Dice (F1) score and Binary Cross Entropy (BCE),

$$\mathcal{L} = \frac{2\sum_{i=1}^{N} p_i g_i + \epsilon}{\sum_{i=1}^{N} p_i + \sum_{i=1}^{N} g_i + \epsilon} - \frac{1}{N}\sum_{i=1}^{N}\left[g_i \log(p_i) + (1-g_i)\log(1-p_i)\right], \quad (3)$$

where the index $i$ runs over all pixels in an image, $p_i$ is the predicted value for that pixel, $g_i$ the groundtruth, and $\epsilon = 10^{-6}$ is a small parameter for stability. The network was trained for 200 epochs on non-augmented data, and then for a further 150 epochs with augmented data.

***Evaluation.*** The performance of the network was evaluated in two ways. First, both the classic method and U-Net were used to segment multiple networks of *R. irregularis*. This is a strain where the classic method works well. The total network length was calculated from the resulting pair of segmentations, and compared in Fig. 7. There is a clear positive correlation between the two measurements, from which we conclude U-Net segmentations are in broad agreement with the classic method. The second evaluation method was to visually compare predictions from U-Net and the classic method made on images with high hyphal density. The classic method is known to perform poorly on such examples (e.g. those from *R. aggregatus*). A comparison is shown in Fig. 8, demonstrating the significant improvement in hypha detection achieved by U-Net in the case of dense networks.

#### High resolution imaging

As described in (Oyarte Galvez et al., 2025). The objective used was 50x instead of 100x but the rest of the system was identical.
We built a customized microscope system to acquire high-magnification videos of cytoplasmic flows inside the mycorrhizal hyphae, with the imaging-path optical system identical to that used for low-magnification network imaging (objective, 200mm tube lens and a Basler acA4112-30um CMOS camera, mounted on a Thorlabs KMTS25E/M motorized stage). However, we used a different objective lens (50X Nikon CFI60 TU Plan Epi ELWD). A 1 W red fiber optic LED light source (Product ID: 4165, Adafruit Industries) for illumination through an LED reflector assembled with a Fresnel lens to collect and diffuse the light before the beam reached the sample. The sample stage was customized X-Y motorized linear stage, with stepper motors (NEMA23 IP20, Servotronix) driven by an Arduino Uno Rev3 micro-controller. With this setup, every video could be related to a specific coordinate in the fungal network.

#### CNN model training and architecture

***Hyperparameter tuning.*** We used keras Bayesian Optimization tuner to explore the hyperparameter space. We set the max number of trials to 20 and the number of initial points to 50. The following parameters were adapted.
*Convolutional Layers*

- Number of Convolutional Layers: Ranges from 1 to 4. The default setting is 2. This determines how many convolutional layers will be added to the model.
- Filters: For each convolutional layer, the number of filters can range from 32 to 256, with a step of 32.
- Kernel Size: The kernel size for each convolutional layer is selectable in a range from 11 to 21 with a step of two. This range was specifically chosen to allow the different hyphal shape to be within the receptive field of the kernel.
- Regularization: Applies L1 regularization on the kernel, activity, and bias with a logarithmic range from 1e-5 to 1e-1 for each, allowing the model to potentially reduce overfitting by penalizing large weights.

*Pooling Layers*

- Pooling: Can be "MaxPooling" (MP), "AveragePooling" (AP), or "No pool". This choice dictates whether to downsample the feature maps and by which method.

- Pooling Size and Padding: For "MP" or "AP", the pooling layer's size ranges from 1 to 4 with step 1, and padding can be either 'valid' or 'same', impacting the downsampling behavior and the spatial dimensions of the output.

*Batch Normalization*

- Inclusion: A boolean indicating whether a Batch Normalization layer is added, aimed at stabilizing and accelerating training by normalizing the inputs of activation functions.

*Dropout*

- Rate: Applied after each dense layer, with a range from 0 to 0.5 in steps of 0.1. This is used to prevent overfitting by randomly setting a fraction of input units to 0 at each update during training.

*Dense Layers*

- Number of Dense Layers: Ranges from 1 to 4, with a default of 2. This defines how many fully connected layers are added towards the end of the model.

- Units in Dense Layers: For each dense layer, the number of units can range from 8 to 256, with a step of 32. This determines the dimensionality of the layer's output space.

**Learning Rate**

- Learning Rate: Used by the Adam optimizer, with a logarithmic range from 1e-5 to 1e-1. The learning rate is crucial for controlling the rate at which model weights are updated during training.

Through Bayesian Optimization, these hyperparameters are systematically explored to identify the combination that results in the best performance on the validation set, as measured by the mean absolute error metric.

***Final architecture.*** Our final model, implemented using the `Keras` framework, is a sequential convolutional neural network designed for processing one-dimensional data. The architecture is summarized as follows:

- The first convolutional layer (`conv_1`) consists of 128 filters with a kernel size of 101 and a stride of 1. It employs the ReLU (Rectified Linear Unit) activation function to introduce non-linearities. This layer is configured with L1 kernel regularization to prevent overfitting by encouraging sparsity in the learned features.

- The second convolutional layer (`conv_2`) also has 128 filters but with a kernel size of 91, maintaining a stride of 1 and using the ReLU activation function. Similar to the first convolutional layer, it includes L1 kernel regularization.

- Batch Normalization: Following the convolutional layers, a Batch Normalization layer is employed to stabilize and accelerate the learning process. This layer normalizes the activations of the previous layer at each batch, maintaining the mean activation close to 0 and the activation standard deviation close to 1.

Dropout Layers:

- The first dropout layer is applied after batch normalization with a dropout rate of 20%. It randomly sets a fraction of input units to 0 at each update during training, which helps in preventing overfitting.

- The second dropout layer follows the first dense layer, with a slightly higher dropout rate of 30%, providing further regularization.

- Flattening Layer: A Flatten layer is used to convert the two-dimensional output of the preceding layers into a one-dimensional array, making it suitable for input into the dense layers. Dense Layers:

- A dense layer with 232 units follows, employing the ReLU activation function. This layer, like the convolutional layers, uses L1 regularization for both kernel and bias. The final output layer has a single unit with a linear activation function, suitable for regression tasks or binary classification.

The total trainable parameters of the model amount to 2,886,097, with an additional 256 non-trainable parameters, leading to a total of 2,886,353 parameters.

***Training.***

**Final Training Procedure** The model was trained using the Adam optimizer with an initial learning rate of 1e-4. The loss function used was mean squared error on the difference between radius squared. We made this choice to avoid underestimation of high radiuses that represented a small proportion of the dataset. The batch size for training was set at 32, and the model was trained for a maximum of 120 epochs. During training, an EarlyStopping callback was employed to prevent overfitting. This callback monitored the validation mean absolute error, with a patience of 20 epochs and a minimum delta of 1e-3 for the first training phase. After the initial training phase, the learning rate was reduced in two subsequent phases to fine-tune the model: In the second phase, the learning rate was set to 1e-5 with the same batch size and number of epochs. The EarlyStopping callback's patience was reduced to 10 epochs. In the third phase, the learning rate was further reduced to 1e-6, retaining the same batch size, number of epochs, and EarlyStopping configuration as in the second phase. This stepwise reduction in the learning rate is a form of learning rate annealing, which helps in fine-tuning the model parameters and potentially improving the model's performance on the validation set.

***Evaluation.***

**Independent Test Set Evaluation** We reserved an independent test set for final evaluation. The model was trained on the entire training set using the final model and learning rates optimized through hyperparameter tuning. It was then used to make predictions on the test set. The performance was assessed using the same RMSE and $R^2$ metrics.

**Comparison with Linear Regression** To benchmark the performance of our neural network model, we compared its results with those from a simple linear regression model. The linear regression model was trained on the same training set and evaluated on the same independent test set. The performance metrics (RMSE and $R^2$) were calculated in the same manner as for the neural network model.

***Results.***

- The null model consisting in taking the average of the training set and using it as a prediction yielded a RMSE of 1.5$\mu$m on the resampled test set.

- The linear regression model, used as a baseline, yielded an RMSE of 0.79$\mu$m and an $R^2$ of 0.58 on the resampled test set.

- On the independent resampled test set, the neural network model achieved an RMSE of 0.70 $\mu$m and an $R^2$ of 0.66.

- The distribution of residuals on the training and test sets show that large radius may tend to be underestimated, and smaller radius overestimated. (SI Fig.1 B,D).

These results indicate that:

- The neural networks does better than the baseline and the simple linear model

- The RMSE of 0.76 on the test set means that the model predicts a radius value with a 95% confidence interval of +-1.5 $\mu$m.

- This error is quite large compared to the typical values of radius. But two points need taking into consideration. When estimating total hyphal surface and volume as it is done in this publication, if radius measurements are (i) independent and (ii) unbiased, error should cancel out. This means that even a large error on radius extraction will translate only in a small relative error on total hyphal surface or biovolume. (i) is justified by the fact that each hyphal radius measurement is taken independently (ii) is not entirely justified given the uneven distribution of residuals, but we can estimate the overall impact of such systematic error.

- The estimated standard deviation of our manual hyphal radius measurement was 0.3$\mu$m. This is about 2 times smaller than the RMSE on the test set, which means the neural network is not so far from the maximum precision it could achieve.

- During prediction, multiple transects are used for each edge, and the median of the predicted radius for these transects is used as the edge radius. This tend to mitigate error at the edge level.

**Estimating integrated error** We estimated the impact of all sources of error (uneven distribution of error along the radius spectrum and quadratic error accumulation) by resampling our radius estimates adding a noise model corresponding to the test set radius dependent RMSE (SI Fig.1D). We specifically divided radiuses in 8 classes from 0 to 8$\mu$m. Then for each edge, we calculated its radius as predicted by the model and added an error sample from the residual distribution for that class (assumed gaussian with mean and standard deviation obtained from the test set data). We followed this procedure on 38 mature networks and found that the total biovolume estimate was affected by a few percent (always inferior to 8%, average 5%). We therefore decided to ignore this source of error in subsequent calculations of 95% confidence interval. The biological variability is indeed of larger amplitude than the uncertainty of biovolume extraction due to imperfect radius extraction.

#### Numerical simulations

The partial differential equations describing traveling-wave growth of the AM fungal network and P depletion were numerically integrated in the same manner as explained in (Oyarte Galvez et al., 2025).

**Flux term** We chose $J(n) = -nv_d\hat{\mathbf{R}} + D\nabla n$ with parameter $D = 0.0018$ mm$^2$/h so that $v_{\text{wave}} = v_d + 2\sqrt{D\alpha} \approx v_d$ and we could more easily vary wave speed.

**Modeling P absorption** Consistent with the P flux used for estimating P absorption by the network, we estimate the local volumetric P depletion in the medium with the equation

$$\phi_P = -\frac{2\pi r\rho J_{\text{max}}[P]}{[P]+K_m} \tag{4}$$

with $K_m$ the Michaelis-Menten constant for absorption. The dynamics of P-concentration in the medium is then determined by the balance of this flux $\phi_P$, which depletes P from the medium, and the diffusive flux $D\nabla^2[P]$, yielding the following partial differential equation (PDE):

$$\frac{\partial[P]}{\partial t} = D_P\nabla^2[P] + \phi_P \tag{5}$$

where $D_P$ is the diffusion coefficient of P in the medium.

**Soil specific P dynamics** As explained in (Oyarte Galvez et al., 2025)P dynamics in agar differ from what can be observed in a real soil. P in soil is in most case reversely bound to soil particles and only a small fraction is in solution. The equilibrium between the liquid fraction concentration $C_L$ and the solid fraction $C_S$ concentration can be represented with the following relations $\frac{dC_S}{dC_L} = b_p$ where $b_p \approx 200$ represents the phosphorous buffer power of the soil (Schnepf and Roose, 2006). Within the regime far from saturation of the solid fraction where $b_p$ is a constant, this simplifies to $C_S = b_pC_L$.

The above equations can therefore be adapted to account for this buffering effect. First the P concentration experienced by transporters at the hyphal surface is $C_L$ and equation. The expression for $\phi_P$ must therefore be adapted to be

$$\phi_P = \frac{sV_{\text{max}}[P]/b_p}{[P]/b_p + K_m} \tag{6}$$

where $[P] = C_S$ represents the total P bound on soil particles. Then neglecting the diffusion of P on solid surfaces, one can rewrite

$$\frac{\partial[P]}{\partial t} = D_P\nabla^2([P]/b_p) - \phi_P \tag{7}$$

This means the diffusion is effectively slowed down by a factor $b_p = 239$ which corresponds to the general consensus for the movement of adsorbed species and specifically Phosphorous in soils (Bieleski, 1973; Darrah and Staunton, 2000). Such value can however vary across soil types with sandy soils having values of $b_P$ closer to $1$ while the value used here corresponds to clayish soils.

**Initial condition** Initial condition were $\rho(R,0) = 0$, $n(R,0) = \frac{q_{max}}{v_g r^2}e^{-\lambda(R-R_0)^2}$ where $q_{max} = 6 \times 10^{-6} mm^3 h^{-1} mm^{-2}$, $\lambda = 1.2mm^{-1}$ and $R_0 = 7mm$ and $[P](r,0) = [P]_0$. The normalization by $v_g$ was used to ensure that all fungal stategies start with an equal increase in volume density representing a fixed initial carbon investment by the plant $r^2\frac{\partial\rho}{\partial t}(R,0) = r^2 v_{\text{dens}} n(R,0) = q_{max}e^{-\lambda(R-R_0)^2}$.

**Other parameters** For all simulations. We chose $\kappa_0 = 3$ corresponding to the value found for AM fungal networks associated with root genotype 1 and $v_g = v_d$. For all simulations, we used $D_P = 3.6mm^2.h^{-1}$ which is the diffusion coefficient of small molecules in water. In Figure 5A, B we choose $k_2 = 0.4mm\,ng^{-1}.h^{-1}$. Figure 5D we choose $k_2 = 0.024mm\,\mu g^{-1}.h^{-1}$. The adaptation rate was choosen differently because in Figure 5A,B we wanted to focus on the long term permanent regime travelling wave dynamics and a too large $k_2$ eventually led to unwanted oscillatory behaviour during integration.

For Figure 4A, B we chose $[P]_0 = 10ng/mm^3$ and varied $v_d$ as shown in the legend and figure axis.

**Integration** In Fig.5A-B: Space was divided in a mesh of 1181 cells from $0$ to $252mm$ and time was divided in 900 elements from $0$ to $T = 900h$. In Fig.5D: Space was divided in a mesh of 525 cells from $0$ to $112mm$ and time was divided in 180 elements from $0$ to $T = 180$.

We ran integration using the `dolfin` libray for integration of PDEs. Code is available in the following repository together with the rest of the replication package.

**Figure 5D** We varied $v_d$ over the interval $[150\mu m/h, 320\mu m/h]$ sampling uniformly 20 times. We varied $[P]_0$ over the interval $[2.5\mu g/mL, 25\mu g/mL]$ sampling 20 times.

### Supplementary discussion

#### Limitations of carbon estimates

- **Variations in Cell Wall Thickness and Lipid Density:** The estimate of carbon flux given in the Methods does not consider variations in cell wall thickness or lipid density inside the cells which can vary from hypha to hypha (Bago et al., 1998, 2002). Further refinements can include such considerations, but we expect the carbon ratio of cells to be close to the one used in this study.

- **Maintenance Cost, Carbon Recycling, and Spores:** At the timescale considered (4-5 days) we considered that some more complex carbon dynamics could be neglected. All the different possible carbon costs for a growing fungal network are nicely summarized and quantified in (Heaton et al., 2015).

  ⋆ First, we neglected the carbon cost associated with cell maintenance. This cost, although of smaller magnitude, is proportional to total network length, while the cost associated with growth is proportional to the derivative of the total length. It has been estimated for fungi that the time over which the metabolic cost of maintaining a living volume is equal to the number of joules embodied in that volume was about 30 days for fungi (Heaton et al., 2015). At the timescale considered it is therefore valid to consider the maintenance cost negligible.

  ⋆ We also neglected the cost associated with transport of resources within the network. This was motivated by the following calculation. Assuming all the cytoplasm is moving at $v_0 = 3\mu m$ in $L = 1m$ of network of radius $R = 3\mu m$. The hydraulic resistance associated with the network is $R_h = \frac{8\mu L}{\pi R^4}$ where $\mu = 1g.m^{-1}.s^{-1}$ is the viscosity of the cytoplasm and the flux through the pipe is $\Phi \approx \pi R^2 v_0$. The energy dissipation power is then of the order of $P_{transport} = R_h \Phi^2 = 8\pi\mu L v_0^2$ which does not depend on $R$. We can therefore evaluate that $P_{transport} = 226 \times 10^{-15} W \approx 0.2 \times 10^{-6} \mu W$. In comparison, we estimate that the growing networks can consume $\phi_C \approx 0.2\mu g/h$ of carbon (see Fig. 2B). This carbon is in the form of lipids/palmitic acid which energy density is about $d_E = 37MJ/kg$. The Power requirement for growth can therefore be estimated to $P_{growth} = d_E \phi_C \approx 2\mu W$. At first approximation, the energetic cost of transport therefore seems negligible compared to the energetic cost of growth.

  ⋆ Fungi can also recycle cell material. To generate the dataset used for machine learning, we sampled multiple positions in the fungal network over several days. At the timescale considered, we didn't observe hyphal cytoplasm retracting except for thin Branched Absorbing Structures (BAS) around the end of that timescale. It was previously estimated that full BAS lifecycle consisted in 7 days for formation followed by 5 weeks until full retraction (Bago et al., 1998). We observe a faster BAS lifecycle in our experiments somewhat (some hyphae starting to retract after just 5 days). Interestingly the timescale of BAS retraction seems to coincide with the one of P depletion. We however decided not to include this effect in our calculation since (i) BAS constitute 30% of total hyphal length (see (Oyarte Galvez et al., 2025)) but are thinner than the rest of the hyphae. We expect they constitute no more than 10-15% of total hyphal volume (ii) While some BAS could indeed start to retract over a similar timescale as the one of our experiments, it was clearly not the case for most of them, so we expect the overall amount of recycling to be negligible at the timescales considered.

- ⋆ Finally, while spores are expected to hold more carbon since they are meant to store carbon, we didn't use spore specific ratios to compute total network carbon. At the timescales considered, spores only constitute a small fraction of total volume. We therefore expect this inaccuracy to be negligible.

- **Carbon investment flux in the root compartment:**
  - ⋆ We're not imaging the extra radical hyphal growth in the root compartment. It is however likely that, because phosphorous is already depleted in that compartment (see Methods), the hyphal growth is also negligible.
  - ⋆ Extra radical hyphal growth is generally accompanied by intra radical hyphal growth that also consumes carbon. However, at the timescales considered (first 10 days), intra-radical growth can be considered negligible in length and probably even more in volume (Oyarte Galvez et al., 2025).
- **Geometrical constraints:** Because our experiments are done in a finite space (half petri dish), it is expected that geometrical constraints could play a role in the allocation of carbon. An extreme case is the fact that when the whole plate is covered with fungal hyphae, no more growth happens and $\Phi_C$ goes to zero. In order to mitigate these effects In the case of Fig. 2 and 4 we:
  - ⋆ Never included data points more than a 100 hour after the network area has reached $200mm^2$
  - ⋆ Never include data points after a maximum time set as the one corresponding to the maximum observed rate of biovolume growth.

#### Limitations of phosphorous estimates

- **BAS and Runner Hyphae:** Estimates of structure specific expression of P absorption genes showed higher relative expression of P-transporters in BAS (Kameoka et al., 2019). The normalisation is however done on volume and not surface, and it is unclear from these results whether the transporter density on the surface of BAS hyphae should be considered higher. We therefore decided not to make any distinction between different hyphal structures beyond their radii.
- **Hyphal Septation:** As explained above, at the timescales considered, we decided not to consider the possibility that some subpart of the hyphal network could retract and septate and therefore not contribute to P absorption. In addition, it is likely that BAS retraction coincides with P depletion and we already account for the fact that hyphae in P depleted region do not contribute to the total absorption by the network.
- **Root compartment absorption:** We only observe hyphae present in the fungal compartment (upper half). However, hyphae that could be absorbing phosphorous can also be present in the root compartment. We do not account for those in our estimates. This is however justified by the fact that at the moment where hyphae are crossing in the fungal compartment, phosphorous is generally already depleted in the root compartment (see Fig.3 and SI section on P measurement).

#### Fitting of Pareto front

In our experiments, growth of the total hyphal length $L$ of the AM fungal network exhibits an initial exponential regime followed by a quadratic travelling wave regime (Oyarte Galvez et al., 2025). In our model, the growth rate in the exponential regime can readily be shown to be proportional to the fixed exchange rate, $\kappa$. Indeed, for a growing network with fixed average hyphal radius $\langle r \rangle$, assuming $[P] >> K_m$ and a fixed exchange rate $\kappa$, we have,

$$\gamma_C \, \pi \xi \langle r \rangle^2 \frac{dL}{dt} = \kappa \, 2\pi r \, J_{\text{max}} \, L, \quad \textbf{(8)}$$

where $\gamma_C$ is the conversion factor between hyphal volume growth rate and the rate of total carbon expenditure by the growing network (including for example spore building). In this initial exponential phase, $\gamma_C$ is constant and $\gamma_C = M_C$. This equation has the solution,

$$L(t) = L_0 \exp\left( \frac{2\kappa J_{\text{max}}}{M_C \xi \langle r \rangle} t \right) = L_0 \exp(\lambda t), \quad \textbf{(9)}$$

where $L_0$ is the initial length, and the growth rate $\lambda \equiv \frac{2 J_{\text{max}}}{M_C \xi \langle r \rangle} \kappa$ is indeed proportional to $\kappa$.
At later times, fungal networkss propagate as a travelling wave characterized by wave speed $v_{\text{wave}}$ and a hyphal length density $\rho_L$ behind the wave front. In this regime, the length evolves quadratically in the half plate:

$$L(t) = \frac{\pi}{2} \rho_L \, v_{\text{wave}}^2 \, t^2. \quad \textbf{(10)}$$

Now if we let $t_0$ denote the time at which the transition between regimes occurs, continuity of $L(t)$ requires

$$L_0 \exp(\lambda t_0) = \frac{\pi}{2} \rho_L v_{\text{wave}}^2 t_0^2. \tag{11}$$

The exchange rate relation also imposes continuity in the derivative of $L(t)$, which yields

$$L_0 \lambda \exp(\lambda t_0) = \pi \rho_L v_{\text{wave}}^2 t_0. \tag{12}$$

Dividing Eq. (12) by Eq. (11) and rearranging, we find

$$t_0 = \frac{M_C \xi \langle r \rangle}{\kappa J_{max}}.$$

Plugging in typical values of paremeters (Table S1) yields $t_0 \approx 50h$.

Substituting $t_0$ back into Eq. (11), we obtain

$$L_0 e^2 = \frac{\pi}{2} \rho_L v_{\text{wave}}^2 \left( \frac{M_C \xi \langle r \rangle}{\kappa J_{max}} \right)^2,$$

which rearranges to

$$\rho_L = \frac{2 L_0 e^2}{\pi v_{\text{wave}}^2} \left( \frac{\kappa J_{\text{max}}}{M_C \xi \langle r \rangle} \right)^2. \tag{13}$$

If we choose $L_0$ so networks of different average radii always start with the same volume $V_0 = \pi \xi \langle r \rangle^2 L_0$ at $t = 0$ (corresponding to a given initial carbon expenditure) and define $\epsilon = \frac{2e^2}{\pi^2}$, we can express this compactly as:

$$\rho_L = \epsilon \frac{V_0}{\xi \langle r \rangle^2} \left( \frac{\kappa J_{\text{max}}}{M_C \xi \langle r \rangle v_{\text{wave}}} \right)^2. \tag{14}$$

Corresponding to the surface area density

$$\rho_S = 2\pi \epsilon \frac{V_0}{\xi^3 \langle r \rangle^3} \left( \frac{\kappa J_{\text{max}}}{M_C v_{\text{wave}}} \right)^2. \tag{15}$$

$V_0$ is an arbitrary parameter that can be adjusted. After computing the Pareto front for each root genotype, we fit Eq. 15 to both fronts using $\kappa = 3$ for genotype 1 and $\kappa = 4$ for genotype 2 using `curve_fit` of `scipy` package of Python. This yields a value for $V_0$ which is then used to plot the curves in Fig. 5C Bottom of Main Text using Eq. 15.

#### Analytical considerations on the optimal wave speed

***Optimal propagation in the non P depleted travelling wave regime.*** Before reaching the permanent regime of traveling-wave growth with P depletion, there is an intermediate regime in which traveling-wave growth is established but the P-depltion front has yet to fully establish. In this regime, when P uptake remains saturated ($[P] >> K_m$), we have

$$\Phi_P(t) = J_{\max} \rho_{\text{L}} \pi v_{\text{wave}}^2 t^2 \, 2\pi \langle r \rangle.$$

Using Eq. 14, we obtain

$$\Phi_P(t) = \frac{1}{\langle r \rangle^3} J_{\max} A \left( \frac{\kappa J_{\text{max}}}{M_C} \right)^2 2t^2 \, \pi^2.$$

Where $A = \frac{\epsilon V_0}{\xi^3}$ This means that strategies with smaller radii will provide higher $\Phi_P$. Given the linear relationship between $v_{\text{wave}}$ and $\langle r \rangle$, this expression clarifies why slower growing strains can achieve better P transfer performance at low $[P]_0$ (Fig. 5D of Main Text).

375 # Supplementary table

**Table 1.** Parameters used across the manuscript

| Parameter | Description | Unit | Value | Used in Fig. | origin |
|---|---|---|---|---|---|
| $d_{\text{cell}}$ | Characteristic cell mass density | $g.cm^{-3}$ | 1.1 | Fig. 2, Fig. 4 and Fig.5 | (Bakken and Olsen, 1983) |
| $f_{\text{dry}}$ | Fraction of dry biomass | . | 0.21 | Fig. 2, Fig. 4 and Fig.5 | (Bakken and Olsen, 1983) |
| $f_{\text{carbon}}$ | Fraction of carbon in dry biomass | . | 0.5 | Fig. 2, Fig. 4 and Fig.5 | (Bar-On et al., 2018) |
| $J_{\text{max}}$ | Maximum uptake rate per unit surface area | ng P $\text{mm}^{-2}\,\text{h}^{-1}$ | 3.3 | Fig. 2, Fig. 4 and Fig.5 | measured |
| $K_m$ | Michaelis-Menten half-saturation constant | µg P $\text{mm}^{-3}$ | $3.1 \times 10^{-5}$ | Fig. 4 and Fig.5 | (Schnepf and Roose, 2006) |
| $p_{respired}$ | Carbon use efficiency during growth | . | 0.5 | Fig. 2, Fig. 4, Fig. 5 | (Manzoni et al., 2012) |
| $\gamma_C$ | Conversion factor between hyphal length biovolume growth rate and carbon expenditure rate | $\mu g P mm^{-3}$ | $231$ | Fig. 5 | $= \frac{d_{\text{cell}} f_{\text{dry}} f_{\text{carbon}}}{1 - p_{respired}}$ |
| $D_P$ | Diffusion coefficent of phosphate ions | $mm^2.h^{-1}$ | 1.8 in Fig. 4D, 3.6 otherwise | Fig. 4, Fig. 5 | (Davison, 2016) |

#### Supplementary figures

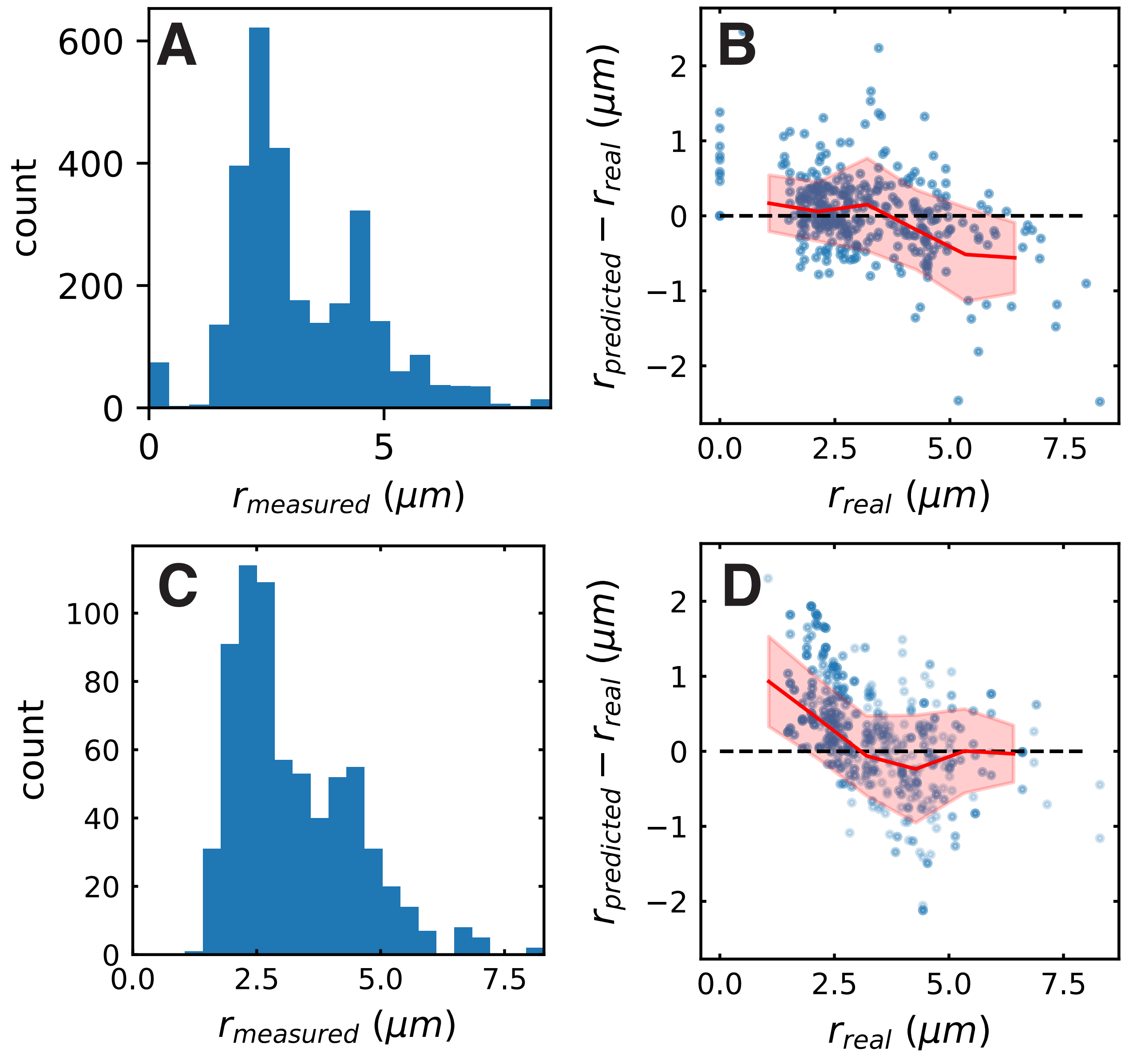


**Figure 1. Training and test set distribution and model performance on validation and test sets.** (**A**) Training + validation set distribution. (**B**) Residual of the model prediction on the validation set. (**C**) Test set distribution after resampling. (**D**) Residuals of the model prediction on the resampled test set. For (**B**) and (**D**), blue points correspond to individual predictions from the set. Radiuses were separated in 8 classes from 0 to 8μm. For each class, the mean residual and the standard deviation were computed. Red line links the mean residual for each class and red shade shows the 95% C.I. for each class (mean ± 2 × s.e.m.).

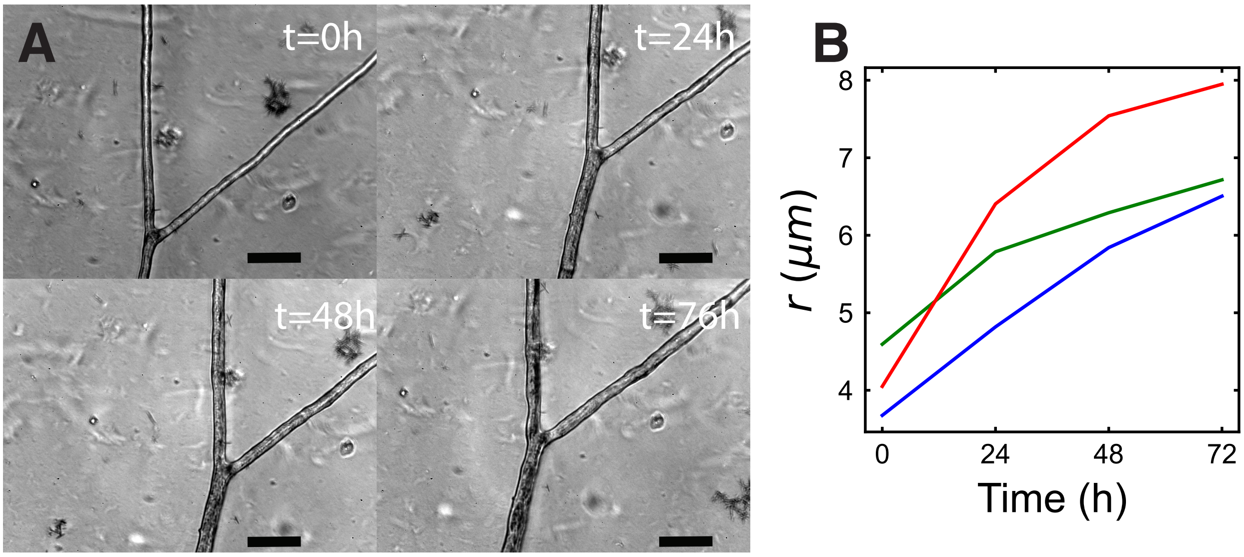


**Figure 2. High resolution imaging shows hyphal widening. (A)** High resolution images of the same Y-shaped intersection over 4 days. Scale bar is 40 $\mu m$. **(B)** Average of measured radius of the three edges of the intersection over time (Bottom : red, upper-left: green, upper-right: blue)

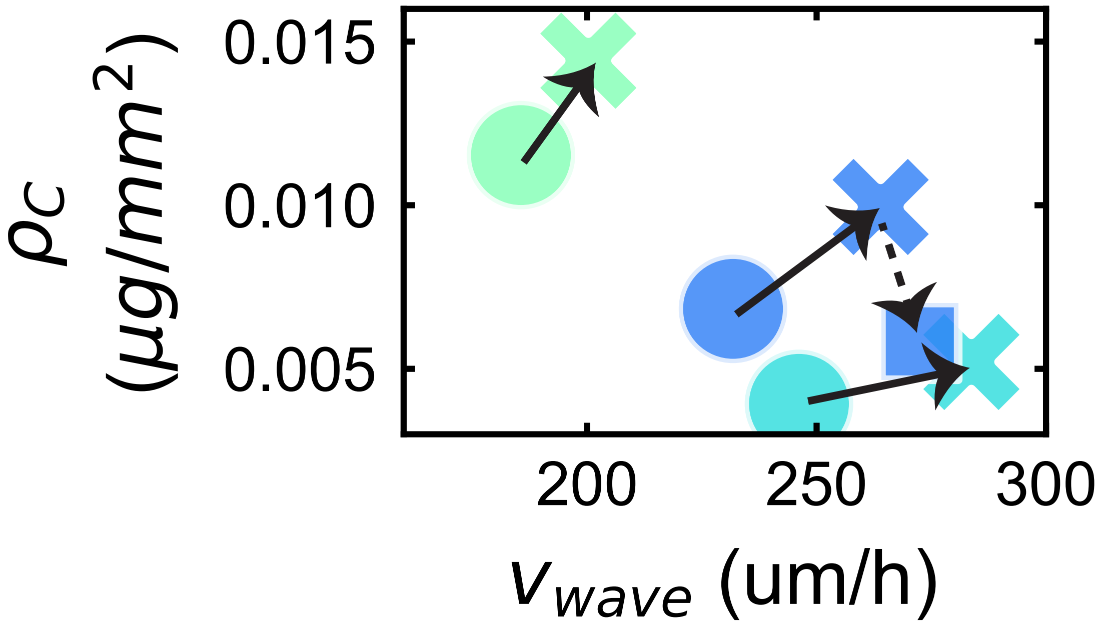


**Figure 3. Effect of changing root host and environment on travelling wave parameters.**Travelling wave observables. $\rho_C$ represents the instantaneous hyphal carbon density and $v_{\text{wave}}$ represents the instantaneous wave speed. Symbols represent the median of those values for each strain. Dark blue corresponds to strain C2, cyan to strain A5 and green to *R. aggregatus*. Circles correspond to carrot root genotype 1 and high P environment in the fungal compartment, crosses correspond to carrot root genotype 2 and high P environment, square correspond to carrot root genotype 2 and low P environment. Arrows are guide for the eye. Continuous arrow corresponds to a change from genotype 1 to genotype 2, dashed arrow corresponds to a change from high to low P environment.

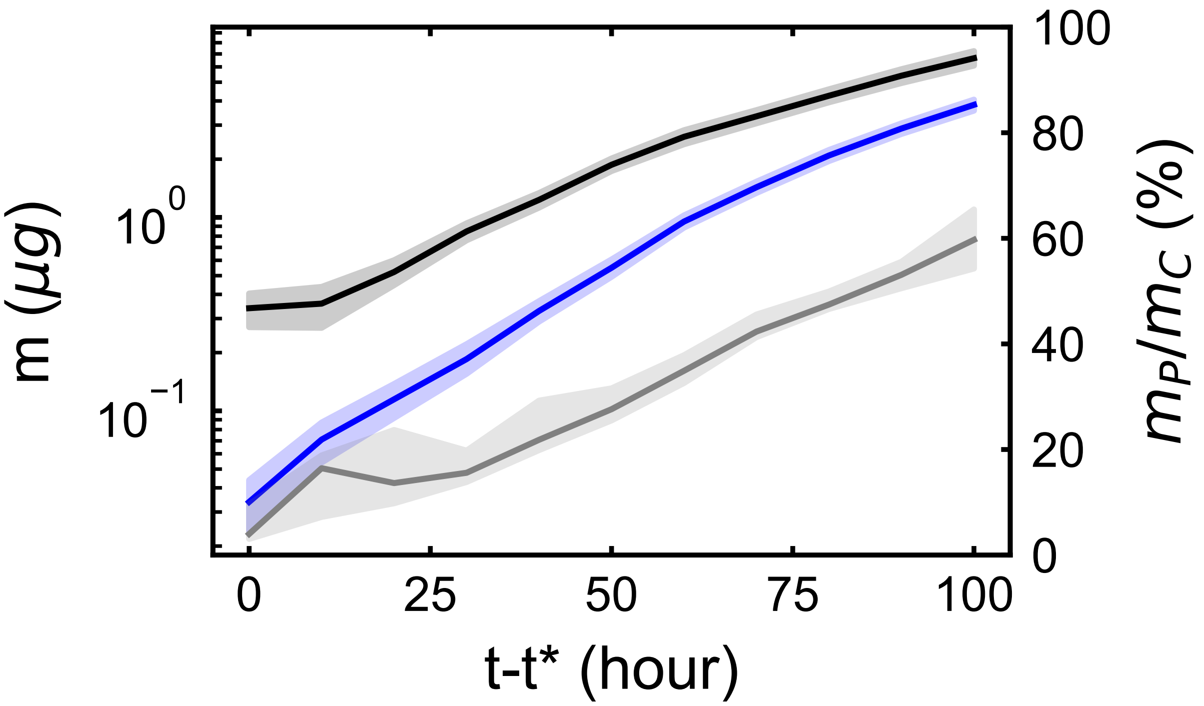


**Figure 4. Mass stoichiometry of fungal hyphae under the hypothesis of no transfer to the plant.** Estimates of total carbon mass (black), total absorbed phosphorous mass (blue) and ratio of the two (grey) as a function of time. Absorbed phosphorous mass is estimated from the measured temporal dynamic of network surface area. Thick lines correspond to average of all replicates over 10 hour time intervals, shades correspond to 95% C.I. over the same interval (mean $\pm 2\times$ s.e.m.).

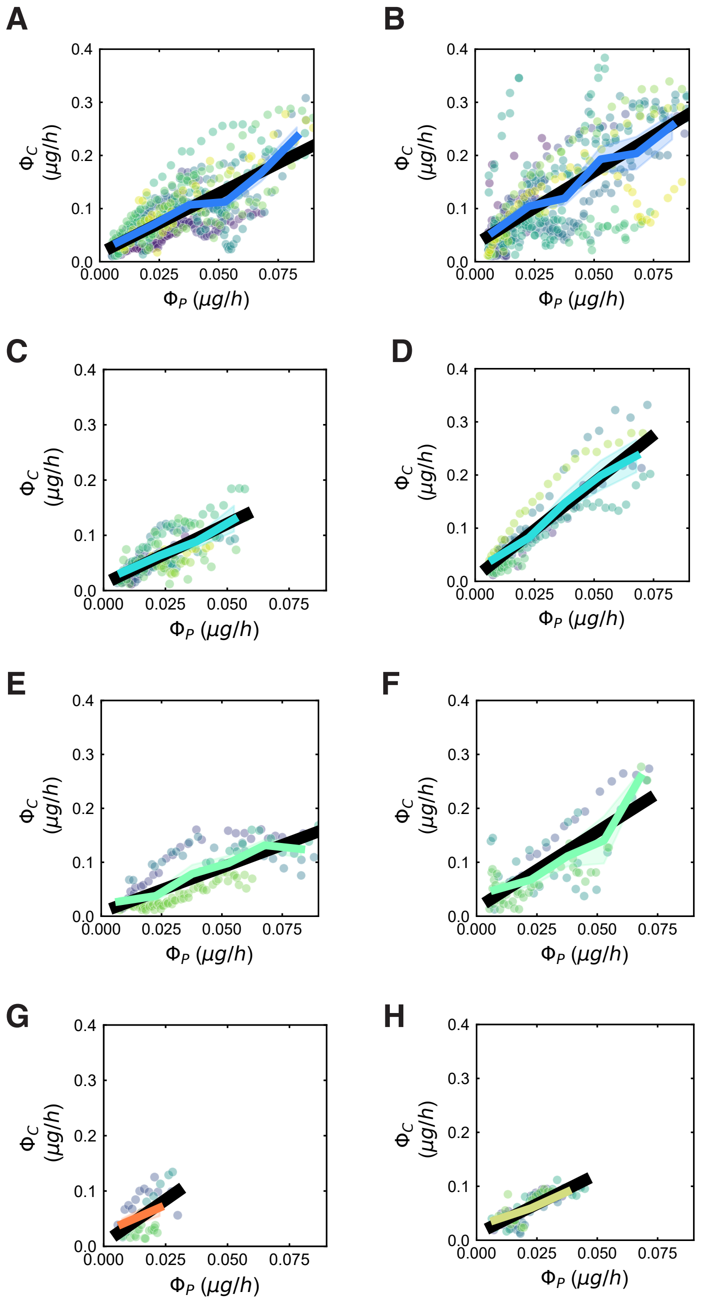


**Figure 5. Proportionality of C expenditure and P supply for each plant hostXfungal strain treatment**Carbon expenditure rate ($\Phi_C$) as a function of Phosphorous supply rate ($\Phi_P$). **(A) ,(C),(E),(G),(H)** correspond to genotype 1 carrot root. **(B) ,(D),(F)** correspond to genotype 2 carrot root. **(A), (B)** correspond to For *R. irregularis* C2. **(A), (B)** correspond to For *R. irregularis* C2. **(C), (D)** correspond to *R. irregularis* A5. **(E), (F)** correspond to *R. aggregatus*. **(G)** corresponds to *R. irregularis* C3. **(H)** corresponds to *G. Clarum* . Each point corresponds to a measurement of $\Phi_C$and $\Phi_P$ at one timestep for one replicate, each replicate is shown in a different colour. Coloured line shade corresponds to binned average 95% C.I. over regular $\Phi_P$intervals. Black line corresponds to linear fit over the blue points. **(B)** For *R. irregularis* A5 (cyan; $n_{\text{genotype 1}} = 8$, $n_{\text{genotype 2}} = 6$ ),C2 (blue; $n_{\text{genotype 1}} = 19$,$n_{\text{genotype 2}} = 11$), *R. aggregatus* (green, $n = 4$), *G. Clarum* (yellow, $n = 4$) associated with genotype 2 (dashed line) and genotype 1 (full line).

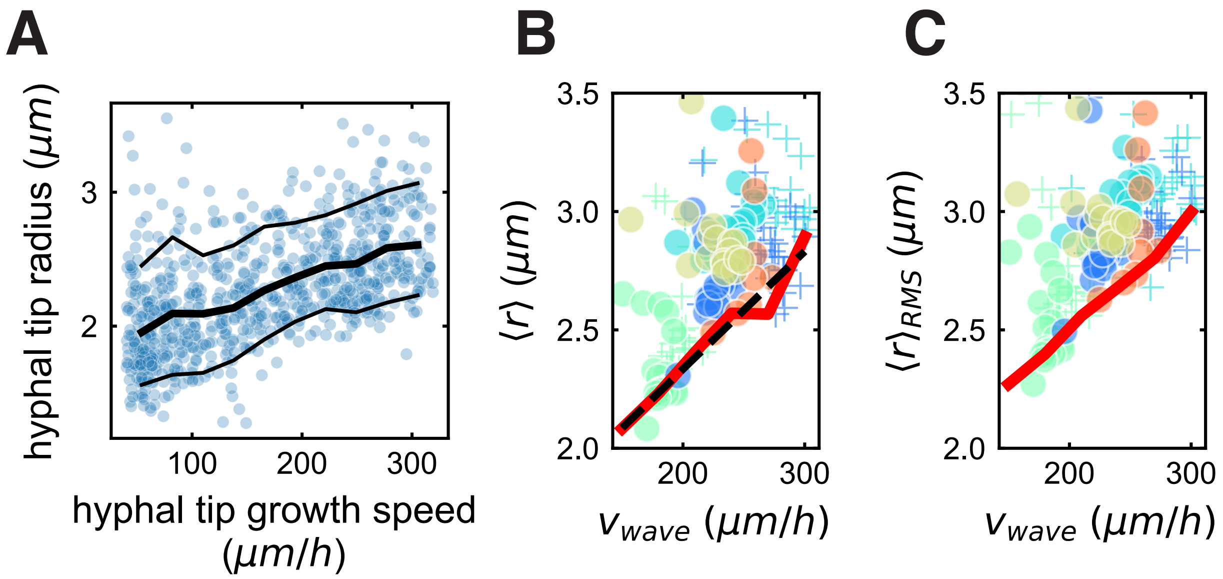


**Figure 6. Fast expansion is constrained by hyphal radius A** Individual hyphal growth speed is constrained by tip radius. Each blue point corresponds to one growth movement of a hypha between two timesteps. Middle thick black line links averages. Upper and lower black lines link respectively the 90th and 10th percentiles. Statistics are computed in 10 bins over the range of hyphal tip growth speeds. **B** Each small circle corresponds to an independent median of mean length weighted radius over 10 hours for all plates of the same strain ($n$ = 7-11). Mean length weighted radius for a given plate at a given time is computed by dividing the newly created total surface area by $2\pi$ times the newly created length. Red line links the minimium over bins of size 30$\mu m/h$). Black line correspond to the linear fit of the points that constitute the red line. **C** Each small circle corresponds to an independent median of length weighted root mean squared radius over 10 hours for all plates of the same strain ($n$ = 7-11). Length weighted root meant squared radius for a given plate at a given time is computed by dividing the newly created total biovolume area by $\pi$ times the newly created length and taking the square toot. Red line links the minimium over bins of size 30$\mu m/h$). In **B** and **C**, colour correspondence and number of replicates are the following: *R. irregularis* A5 (cyan, n=8), C2 (blue, n =19) and C3 (red, n=2), *R. aggregatus* (green, n=4), *G. Clarum* (yellow, n= 4) .

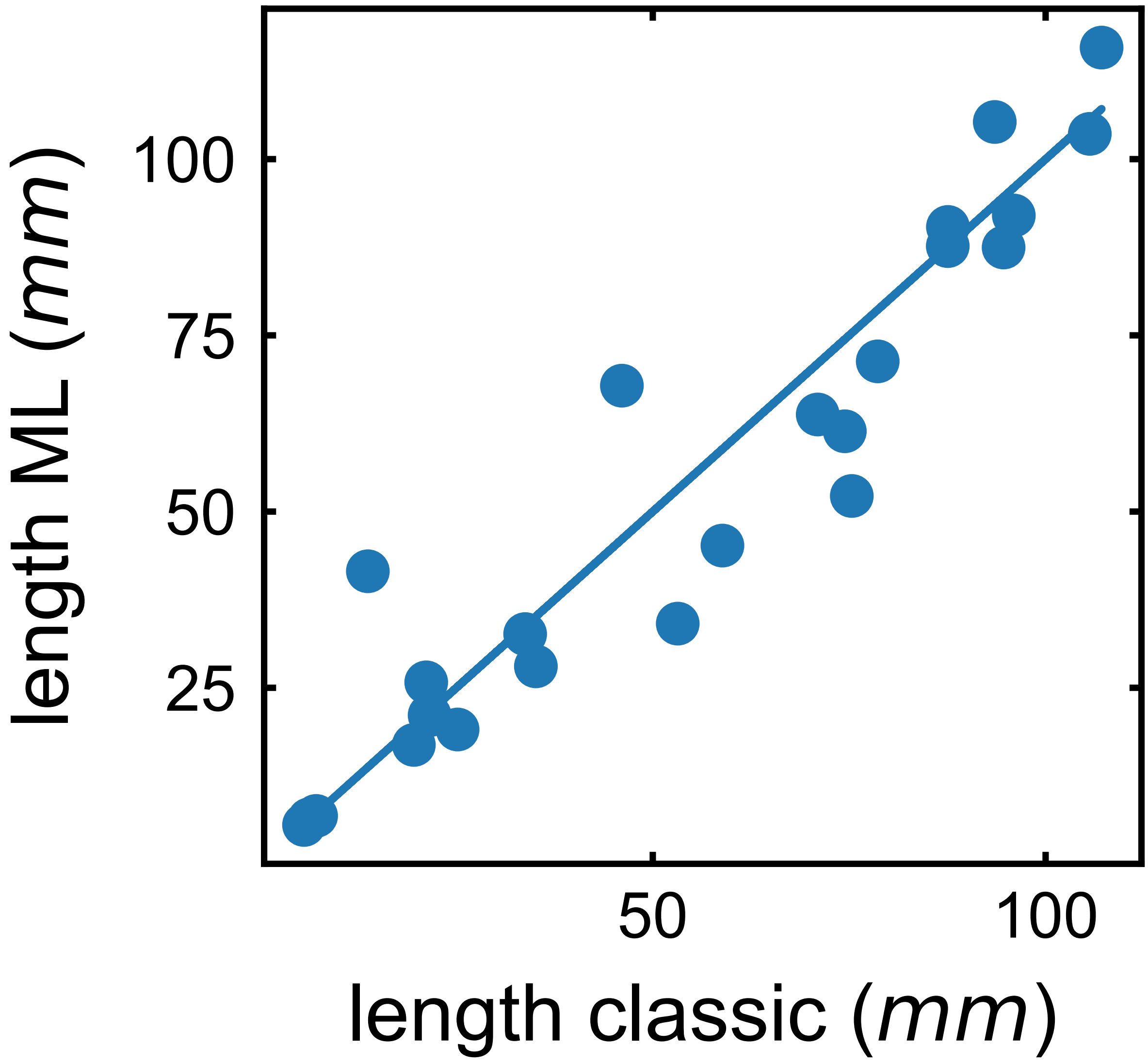


**Figure 7. Two segmentation methods yield similar results on *R. irregularis*.** Each point is obtained from comparing total length obtained through segmentation of a single tile either with the classic algorithm or with the machine learning based one. Blue line is a 1:1 line.

.

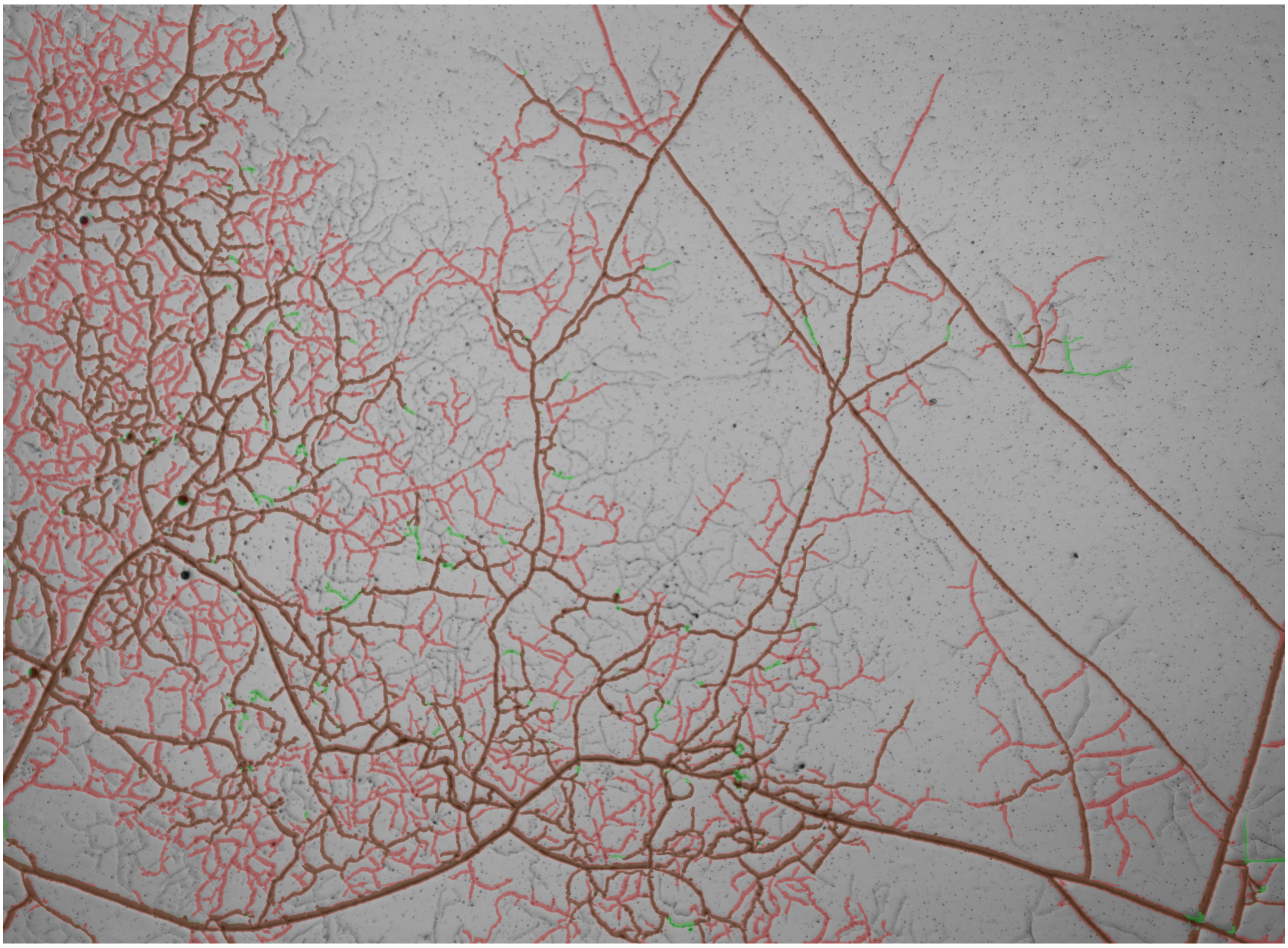

**Figure 8. U-Net segmentation captures many more hyphae in dense networks.** The output of U-Net is shown in red, the classic segmentation method is in green, and brown regions are where both colors overlap.
